# Modularity and connectivity of nest structure scale with colony size

**DOI:** 10.1101/2021.04.30.442199

**Authors:** Julie S. Miller, Emma Wan, Sean O’Fallon, Noa Pinter-Wollman

## Abstract

Large body sizes have evolved structures to facilitate resource transport. Like unitary organisms, social insect colonies must transport information and resources. Colonies with more individuals may experience transport challenges similar to large-bodied organisms. In ant colonies, transport occurs in the nest, which may consist of structures that facilitate movement. We examine three attributes of nests that might have evolved to mitigate transport challenges related to colony size: (1) subdivision - nests of species with large colonies are more subdivided to reduce crowd viscosity; (2) branching - nest tunnels increase branching in species with large colonies to reduce travel distances; and (3) short-cuts – nests of species with large colonies have cross-linking tunnels to connect distant parts of the nest and create alternative routes. We test these hypotheses by comparing nest structures of species with different colony sizes in phylogenetically controlled meta-analyses. Our findings support the hypothesis that nest subdivision and branching evolved to mitigate transport challenges related to colony size. Nests of species with large colonies contain more chambers and branching tunnels. The similarity in how ant nests and bodies of unitary organisms have evolved in response to increasing size suggests common solutions across taxa and levels of biological organization.

## INTRODUCTION

The consequences of body size for the evolution of phenotypic diversity are far reaching, influencing morphological (Schmidt-Nielson 1984), physiological (Kleiber 1947), and behavioral traits (Calder 1984; Dial et al. 2008) in systematic ways (Peters 1983). Evolutionary allometry (Thompson 1917; Huxley 1932), the study of how body size relates to trait size across lineages (Gould 1971; Cheverud 1982), has offered a window into the underlying mechanisms that drive trait diversity (e.g. McMahon 1975; Fairbairn 1997; Frankino et al. 2005). An important consequence of increasing body size is that it alters the basic physical conditions in an organism by increasing the distances over which transport of nutrients, waste products, and signals must traverse (Bonner 2004). This physical transport challenge has been resolved through the evolution of a variety of structural features across the tree of life (McMahon and Bonner 1983), including multi-cellularity (Niklas and Newman 2016), body plans that increase surface-area-to-volume ratios (e.g., jellyfish and sponges; Rupppert et al. 2004), branching vascular systems in animals and plants (West et al. 1997; Banavar et al. 1999), and compartmentalization of specialized tissues into organs (Gould 1966).

Social insect colonies are often compared to whole organisms (Wheeler 1910; Seeley 1989; Holldobler and Wilson 2009), and large colonies may experience transport challenges that are similar to those of large bodied organisms (Gillooly et al. 2010). In larger colonies, the speed and/or reliability with which resources, or information, reach disparate parts of the colony will depend on transfer occurring among more individuals and across greater distances (Pacala et al. 1996; O’Donnell and Bulova 2007; Dornhaus et al. 2012). The outstanding diversity of the ant lineage (Hölldobler and Wilson 1990), and multiple independent evolutionary increases in colony size (Burchill and Moreau 2016), provide an ideal foundation for comparative work that investigates how physical structures evolved to mitigate transport challenges related to size.

Transport of resources and information in social insect colonies may depend on the layout of their nests. Nest structure affects the routes that individuals take as they move resources and information. Therefore, nest structure influences the way in which individuals interact (Pinter-Wollman et al. 2013; Pinter-Wollman 2015b; Tschinkel 2015; Pinter-wollman et al. 2018; Vaes et al. 2020) and behave as a collective (Burd et al. 2010; Pinter-Wollman 2015a). Having more individuals may slow the delivery of resources and information, but the physical structure of the nest can promote rapid movement. Here we investigate macroevolutionary trends in how key nest traits, including subdivision, branching, and short-cuts, relate to typical colony size, i.e., the average number of individuals in a colony, in ants (Formicidae).

Subdivision, or compartmentalization, is one structural feature that large systems evolved to mitigate the challenges of size. For example, large multicellular organisms consist of subdivided cells and organs (Kempes et al. 2012; Okie 2013). However, the size of the units are constrained by the costs of expanding surface area-to-volume ratios (Bonner 1988; Lorenz et al. 2011). In social insect nests, larger nest chambers can hold more individuals, but make it harder for individuals to contact each other (Brian 1956; Aguilar et al. 2018). Constraining the size of nest chambers reduces the number of individuals occupying each chamber while reducing the distances they need to travel to interact. The “subdivision hypothesis” predicts that the nests of species with larger colony sizes will consist of more chambers compared to nests of species with small colonies, while chamber sizes will not differ in relation to colony size.

Another way that large systems have evolved to improve transport is by increasing their connectivity and in turn, reducing travel distances. Travel distances can be reduced by adding paths that branch off from main routes. Many natural systems exhibit branching structures which reduce travel distances without the addition of cross-linking paths. For example, hierarchical branching structures are common in the vascular networks of animals (West et al. 1997; Tekin et al. 2016) and plants (McKown et al. 2010; Blonder et al. 2018). Branching structures are common when resources enter through a single site and need to be homogenously distributed throughout the system (Dodds 2010). The “branching hypothesis” predicts that nests of species with larger colonies will have tunnels with high branching frequency to reduce travel distances in the nest.

Reducing travel distances and increasing connectivity can further be accomplished by adding cross-linking paths, or short-cuts, that connect distant parts of a network and create alternative access routes to maintain traffic flow (Corson 2010; brains: Bullmore and Sporns 2012; plants: Katifori et al. 2010; Blonder et al. 2018; termite nests: Valverde et al. 2009). The “short-cut hypothesis” predicts that nests of species with larger colonies will contain more cross-linking paths than nests of species with smaller colonies. Despite the transport benefits of adding short-cuts, every additional connection has a building and maintenance cost, and highly connected nests may be structurally unsound (Monaenkova et al. 2015).

Ants offer an ideal system to study how nest structure co-evolves with group size on macroevolutionary scales because ants span a large range of colony sizes (Bourke and Franks 1995), and exhibit tremendous structural diversity in their nests (Tschinkel 2015). Early lineages of ant species have small to medium sized colonies (Burchill and Moreau 2016) and build small nests with a single tunnel and/or chambers (Tschinkel 2003), similar to their Apoid wasp ancestors (Branstetter et al. 2017). The evolutionary radiation of the ants resulted in a diversity of nest structures and forms, but their diversity has not been quantified and related to patterns of colony size evolution. Furthermore, it is unknown how the transport challenges introduced by large colony sizes have been mitigated in the evolutionary history of ants. Here, we focus on ground-nesting species because, out of all ants, they have the greatest potential to control the structure of their nests.

Prior studies of subterranean ant nest structure in relation to colony size have focused on the ontogeny of single species, showing that ant nests grow in volume and increase chamber size and/or number as colonies age (Tschinkel 1987, 2004, 2005, 2011, 2015). Fewer studies have examined how tunnel connectivity changes with colony development (Buhl et al. 2004a), so branching patterns in ant nests are not well understood. Furthermore, these studies on the ontogeny of nest structure do not provide information on the evolution of nest structures in relation to mature colony sizes, which we examine here in a phylogenetically controlled cross-species comparison.

Here we investigate interspecific variation in ant nest structure and colony size within a phylogenetic context. We consider an adaptive hypothesis for the evolution of nest structure diversity: if ant colonies face size-associated challenges in transport and communication, analogous to other systems, then the structure of nests should relate to colony size in ways that ease the movement of individuals, resources, and information throughout the nest. We quantify three attributes of nest structure, (1) subdivision, (2) branching, and (3) short-cuts, and examine whether these traits relate to colony size in phylogenetically controlled, cross-species meta-analyses. We further consider that variation in nest structure could be the result of non-adaptive processes, like genetic drift or constraints imposed by evolutionary history. To determine the extent to which phylogenetic history predicts variation in nest structure, we measure the phylogenetic signal for the above nest traits and colony size.

## METHODS

### Obtaining nest and colony size data

To test our hypotheses, we gathered data on nest architecture of subterranean ant species. We identified publications with graphical representations of ant nests by searching for the terms “nest”, “structure”, “architecture”, “casting”, and “excavation/ed” on Google Scholar and Web of Science. We identified 24 papers that contained photographs or illustrations of castings or excavations of complete ant nests with distinguishable entrances, tunnels, and chambers. For the subdivision hypothesis, we included data from 3 additional papers which contain tables with measures of chamber size and number, even though nest images were not available (Table S1). We excluded 14 publications from our dataset because interior chambers and/or tunnels were obscured, nests were constructed under artificial conditions, or were from incipient colonies (Table S2). Authors were contacted for additional images when parts of the nest were obscured, accounting for approximately 10% of the images we used. Physical castings of unpublished nests were also obtained (Table S1). When more than one nest was available for a species, we took an average of all measurable nests.

To relate nest structure with colony size – i.e., the number of ants in a colony - we searched the scientific literature and the web (AntWiki.org and AntWeb.org) for information on the number of workers in a mature colony. For the majority of species, we used the average colony size from the literature. When only a range (minimum - maximum) for number of ants was available, we took the midpoint. Information about the number of ants in a colony at the time of nest excavation or casting was available in less than half of the studies we obtained nest structures from, but when available, we used the reported number of adult workers for those particular nests. When colony size was not available for a particular species, or when nest identity at the species level was unknown, we used colony sizes representative of the order of magnitude for the genus obtained from a sister species. This approximation was necessary for the *Forelius sp* nest cast in Anza Borrego, CA (Booher, personal communication). At this site, only two species of *Forelius* occur and they have the same reported colony size, so we used the colony size from one of them - *F. pruinosus* (10^4^). For *Dorymyrmex bureni* we used the only species from this genus with a recorded colony size - *Dorymyrmex bicornis* (10^3^) (Table S1). We excluded *Pheidole oxyops* and *Acromyrmex subterraneus* nests from our analysis because colony size was not available for these species, and because we could not make a reliable approximation due to substantial variation in recorded colony sizes (Table S2).

Nests in our sample come from species belonging to 6 subfamilies, including from the most diverse subfamilies (Myrmecinae, Formicinae, and Dolichoderinae), and the early branching Ponerinae, which tend be characterized by smaller colony sizes and more basal traits in general. However, we lacked nest data from other early lineages, like the Leptanillinae and the Amblyoponinae.

### Nest Subdivision Hypothesis

To test the subdivision hypothesis, we related colony size to the number and size of nest chambers. We counted chambers in 296 nests from 43 species in 24 genera and measured their sizes in 188 nests from 37 species in 21 genera. To quantify the size of chambers, we measured the maximum width of each chamber using ImageJ (Rueden et al. 2017). We excluded nests from the size analysis if images did not include a scale bar or if species identity was unknown for a nest (e.g., *Forelius sp.*). We used chamber width as a proxy for chamber area because we relied on 2D images of 3D structures. When we had access to physical castings, we measured chamber width directly from the structures using a measuring-tape. We validated that chamber width was a reliable indicator of chamber area using an independent dataset (see *Chamber Width Validation* section of the Supplemental Materials).

To allow for cross-species comparison, we standardized chamber width to worker size. Worker size was measured using a standard metric of ant body size, Weber’s length (Brown 1953), using ant profile images containing a scale bar from AntWeb.org or other publications. For all species, we measured the available images up to 8 specimens (1-8). For monomorphic species and for species with continuous worker polymorphism, we took the average length from all available images. For species with discrete worker polymorphism (i.e., majors and minors), we used a weighted average, based on typical frequency of majors according to the literature (Brown and Traniello 1998; Murdock and Tschinkel 2015). Standardized widths of all chambers in each nest were averaged to obtain a single standardized chamber width per nest.

### Nest Connectivity

#### Nests as Networks

To examine nest connectivity, we depicted each nest as a network, as in (Buhl et al. 2004b; Perna et al. 2008; Viana et al. 2013; Pinter-Wollman 2015a; Kwapich et al. 2018), by manually assigning a node ID to each chamber, nest entrance, tunnel junction, and ends of tunnels that did not reach a chamber (‘end nodes’) and connecting these nodes with edges depicting tunnels. See Figure S1 for an illustration of this method. When quantifying nest casts as networks we excluded the "mushroom top" where the casting material pools around the nest entrance when it is poured (Tschinkel 2010). We designated the nest entrance as the narrowest section of the mushroom top. We obtained networks of 170 nests of 38 species from 21 genera, excluding images in which the network structure could not be inferred because tunnel connections were obscured or ambiguous (Table S1 & 2). We excluded polydomous nests because it was not possible to calculate the network measures we used, mean travel distances and number cycles (see below), without information about the structure of trails connecting different nests.

#### Branching Hypothesis

To test the if branching reduces travel distances in large colonies, i.e., the ‘branching hypothesis’ we quantified the length of the travel paths in a nest as the mean distance of a network. Mean distance of a network is the sum of the lengths of the shortest paths that connect all possible pairs of nodes, divided by the number of all possible node pairs (Figure 1):

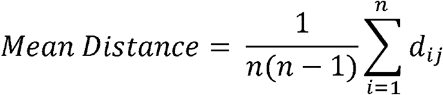

where *n* is the number of nodes, and *d* is the shortest path between all node pairs *i* and *j* (Newman 2018). We used the “mean_distance()” function in igraph R package (Csárdi 2020).

**Figure 1:**
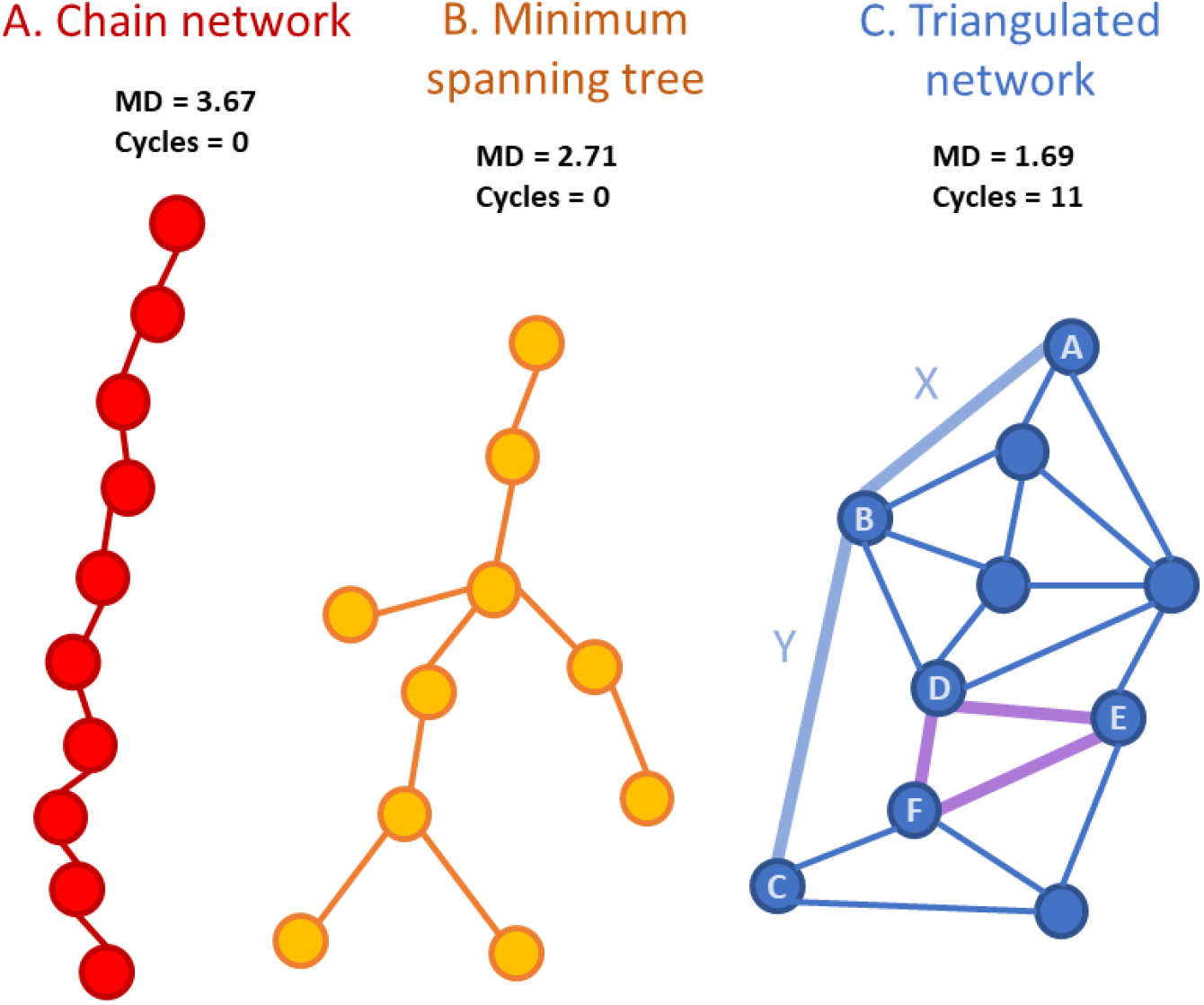
Examples of nest topologies with 10 chambers connected based on the three generative reference models we tested: (A) Chain network, (B) Minimum Spanning Tree, and (C) Fully connected Triangulated network. Above each network we provide its mean distance (MD), which is the average of shortest paths that connect all possible pairs of nodes. As an example, in (C), the length of the shortest path between nodes A and B is one, indicated by the light blue edge X. The length of shortest path between nodes A and C is two, indicated by the light blue edges X and Y. We further provide the number of cycles in each network. As an example, in (C) the purple edges are a cycle connecting nodes D, E, and F.

To determine if observed nests had more branching than expected based on their size, i.e., the number of chambers, or nodes, in the network, we compared the mean distance of networks from observed nests to mean distance of reference models of networks with known branching properties. We evaluated how mean distance relates to nest size under different assumptions about how new nodes are added to a network (i.e., generative models (Hobson et al. 2021)). We used three generative reference models that represent the upper and lower bounds on connectivity for how nest networks might increase in size (Buhl et al. 2004b; Bebber et al. 2007; Tero et al. 2010; Latty et al. 2011): (1) chain networks (Figure 1A), in which new nodes are added to the last node in a chain – i.e., no branching; (2) triangulated networks (Figure 1C), in which new nodes are connected to at least two other nodes such that they form a triangle while avoiding edge overlap. We generated random triangulated networks using Delaunay triangulation with the deldir() function in the ‘deldir’ R package (Turner 2020). The x,y coordinates of the nodes to connect were randomly generated using the ‘sample()’ function; and (3) minimum spanning trees (MSTs; Figure 1B), in which each new node is added to a terminal node to form a new branch in a hierarchical structure. New nodes are added such that the total length of the network is minimized and no cycles (Figure 1C) are formed. We generated MST networks by pruning the triangulated graphs generated in (2) using the ‘mst’ function in the ‘igraph’ R package (Csárdi 2020). Each of the above generative processes was run for network sizes ranging from 3 to 100 nodes, in increments of one node. We repeated this process 1000 times for each network size for both triangulated networks and MSTs, because they were probabilistic generative processes, but not for the chain network generation because each size has only one solution. For each generated network, we calculated the mean distance to create reference distributions for comparison with the observed data (Hobson et al. 2021; Figure 3B). See R code in the supplementary materials.

We further scaled the observed networks to the reference models to relate mean distance to colony size (number of individuals) in a way that accounted for expected changes in network size (number of nodes). To create scaled measures of mean distance, we divided the observed mean distance by the mean value of mean distance of the reference networks (from the 1000 simulations) for each network size (3-100 nodes). The closer the scaled mean distance value is to 1, the more similar the observed network is to the reference model. Scaled values greater than 1 indicate that the observed network has a higher mean distance than expected according to the reference model. Likewise, scaled values less than 1 indicate a smaller mean distance than expected according to the reference model. We related these scaled values with colony size using phylogenetically controlled comparisons as described below in Statistical Analyses (Figure 3C).

#### Short-Cut Hypothesis

To test if larger colonies have nests with more short-cuts, i.e., the short-cut hypothesis, we quantified short cuts using the number of cycles in a network (Buhl et al. 2004b; Pinter-Wollman 2015a). A cycle is a path along a network that returns to its starting node after passing through at least one other node (Newman 2018; Figure 1C). The number of cycles in a nest is:

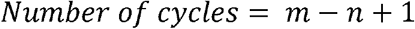

where *n* is the number of nodes and *m* is the number of edges. Cycles are a good approximation of path redundancy of a network, or presence of short-cuts, indicating how often there is more than one route to any given part of the nest (Perna and Latty 2014).

To account for how network size affects the rate at which new cycles are added, we compared the observed nests to the reference models of networks described above. Following a similar procedure to the one described for network mean distance, we calculated the number of cycles in triangulated networks (Figure 3E) and then scaled the number of cycles in observed nests to triangulated networks of equivalent size. Because both chain and MST reference models do not produce cycles, we did not include these models when we scaled the observed nests. We related the scaled number of cycles in a nest with colony size using phylogenetically controlled comparisons as described below (Figure 3F).

### Statistical Analyses

We examined the effect of colony size on the above nest measures while controlling for phylogeny. We used a pruned evolutionary tree of the ants (Formicidae) from (Blanchard and Moreau 2017) and calculated the mean trait value for each species. A complete species-level tree was not available for all species in our dataset, so we selected only one species per genus and ran our analysis using a genus-level tree (Figure 4). When there were data for more than one species per genus, we used the species which had the most nests in our dataset (subset 1 in Tables S3-S13). To ensure this sub-setting of species did not bias our results, we re-ran each analysis with previously excluded species as representatives of a genus, and report the proportion of those runs that were statistically significant. See supplementary materials for results from all runs (Tables S3-13).

We ran phylogenetic generalized least squares (PGLS) comparing each nest trait to the log of colony size. PGLS is a multiple regression that controls for the degree of similarity between species based on phylogenetic distance in a co-variance matrix. We ran PGLS by estimating the phylogenetic signal, lambda λ, through maximum likelihood in the “caper” package v1.0.1 in R (Orme et al. 2018), with λ=0 meaning no phylogenetic signal, and λ=1 meaning the observations completely match phylogeny (Pagel 1999). Due to the small sample size (20 - 24 genera) we could not estimate lambda with high confidence (Freckleton 2009). Therefore, we further conducted PGLS for fixed lambda values (0, 1) using the “ape” package v5.1 in R (Paradis et al. 2018). When our models did not meet the assumption of normality of residuals we analyzed these relationships by calculating phylogenetic independent contrasts (PICs) instead (Paradis et al. 2018). PIC transformed data were modeled using OLS regression, with the intercept set to the origin (R Core Team 2019). Due to small sample sizes, we calculated 95% confidence intervals for OLS model coefficients using a bootstrap (1000 iterations).

To estimate phylogenetic signal for all our measures of nest structure and the log of colony size, we estimated Pagel’s lambda with the ‘phylosig’ function in the “phytools” R package (Revell 2017). This function estimates the value of Pagel’s lambda from 1000 simulations and compares it to a model of no phylogenetic signal (λ=0) using a likelihood ratio test. Due to small sample size, we also compared the fit of our estimated lambda model of phylogenetic signal to a Brownian Motion model (λ=1) and to a model with no phylogenetic signal (λ=0) using AICc values calculated from the ‘fitContinuous’ function in the “geiger” package in R (Harmon et al. 2020). Finally, to visualize colony size and nest trait variation across the phylogeny, we mapped standardized trait values onto a genus level phylogeny from species subset 1 (Figure 4) using the ‘contMap’ and ‘phylo.heatmap’ functions in the “phytools” R package (Revell 2017).

## RESULTS

### Subdivision Hypothesis

Both predictions of the subdivision hypothesis, that chamber number would be larger in species with large colony sizes and that chamber size would not relate to colony size, were supported. We found a statistically significant positive relationship between number of chambers per nest and colony size (Figure 2A; N= 24 genera, OLS of PIC: β = 4.024, 95% CI = 2.014, 6.292, R^2^ = 0.560, *p* < 0.0001; Table S3). The effect size of this relationship was large (3.7 < β < 4.3 across all data subsets) and statistically significant in all species subsets (19/19 datasets). We did not detect a statistically significant relationship between the average standardized chamber width and colony size (Figure 2B; N=21 genera, OLS of PICs: β = 0.350, 95% CI = −0.049, 0.959, R^2^ = 0.140, p = 0.095; Table S4) in the majority of the species subsets (13/15 datasets).

**Figure 2:**
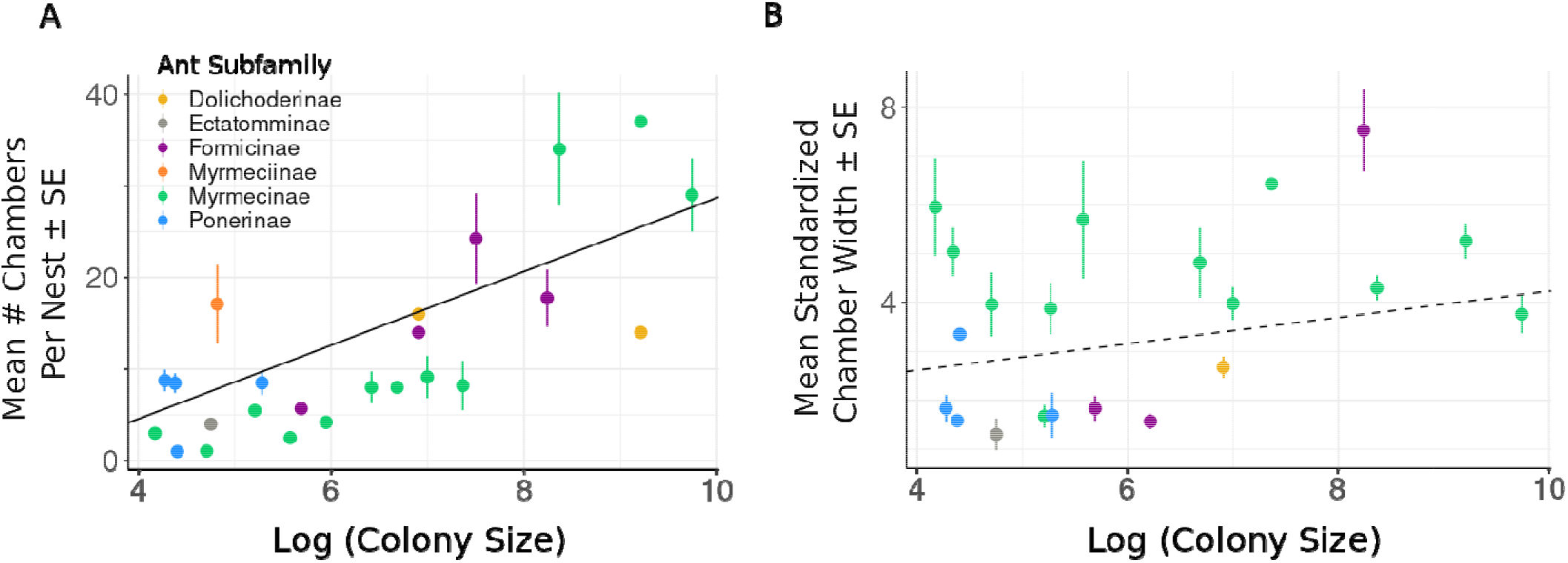
Relationship between the log of colony size (number of ants in a colony) and (A) average number of chambers per nest or (B) average standardized chamber width. Each data point represents a mean value for a single species, whiskers indicate standard errors when multiple nests were sampled from the species. Colors represent ant subfamilies. Lines are fits from the PGLS (λ=1) with a statistically significant fit as a solid line and a relationship that is not statistically significant as a dashed line.

### Branching Hypothesis

The “branching hypothesis”, that structural branching would reduce travel distances in nests of species with large colonies, was supported. Raw, unscaled, mean distance of nests significantly increased with colony size (Figure 3A; N = 21 genera, PGLS λ=MaxLL: β =1.36 ± 0.247, λ=0.585, R^2^ = 0.615, *p* = 0.0007; λ=1: β = 0.844 ± 0.227, p=0.0015; λ=0: β = 1.013 ± 0.251, p<0.0001; Tables S5-S7). In the chain network reference models, mean distances increased fastest and linearly with network size (i.e., number of nodes; Figure 3B). In the triangulated reference models, mean distances were smallest and increased nonlinearly, slowing their rate of increase with increasing network size. In the MST reference models, mean distances were intermediate between the values expected from the chain and triangulated reference models, and also increased nonlinearly (Figure 3B). The mean distances of empirically observed nests were between the expected values for chain and MST reference models (Figure 3B). When we scaled the observed mean distance with the expected mean distance based on each of the three reference models, we found that the relationship between scaled mean distance and colony size differed across the three models (Figure 3C). Mean distances scaled to chain networks declined with colony size and the relationship was statistically significant in almost all data subsets (15/16 datasets; PGLS λ=1: β = −0.055± 0.024, *p* = 0.031; 16/16 datasets; λ=0: β = −0.061 ± 0.024, *p* = 0.018; Tables S8 & S9). In contrast, mean distances scaled to triangulated networks increased significantly with colony size in almost all the data subsets (15/16 datasets; OLS of PIC: β = 0.236, 95% CI = 0.055, 0.466, R^2^ = 0.348, *p* = 0.0048; Table S10). Mean distances scaled to MSTs also increased significantly with colony size (15/16 species subsets), but the effect size was smaller than for triangulated networks (OLS of PIC: β = 0.11, 95% CI = −0.022, 0.240, R^2^ = 0.281, *p* = 0.013; Table S11).

**Figure 3:**
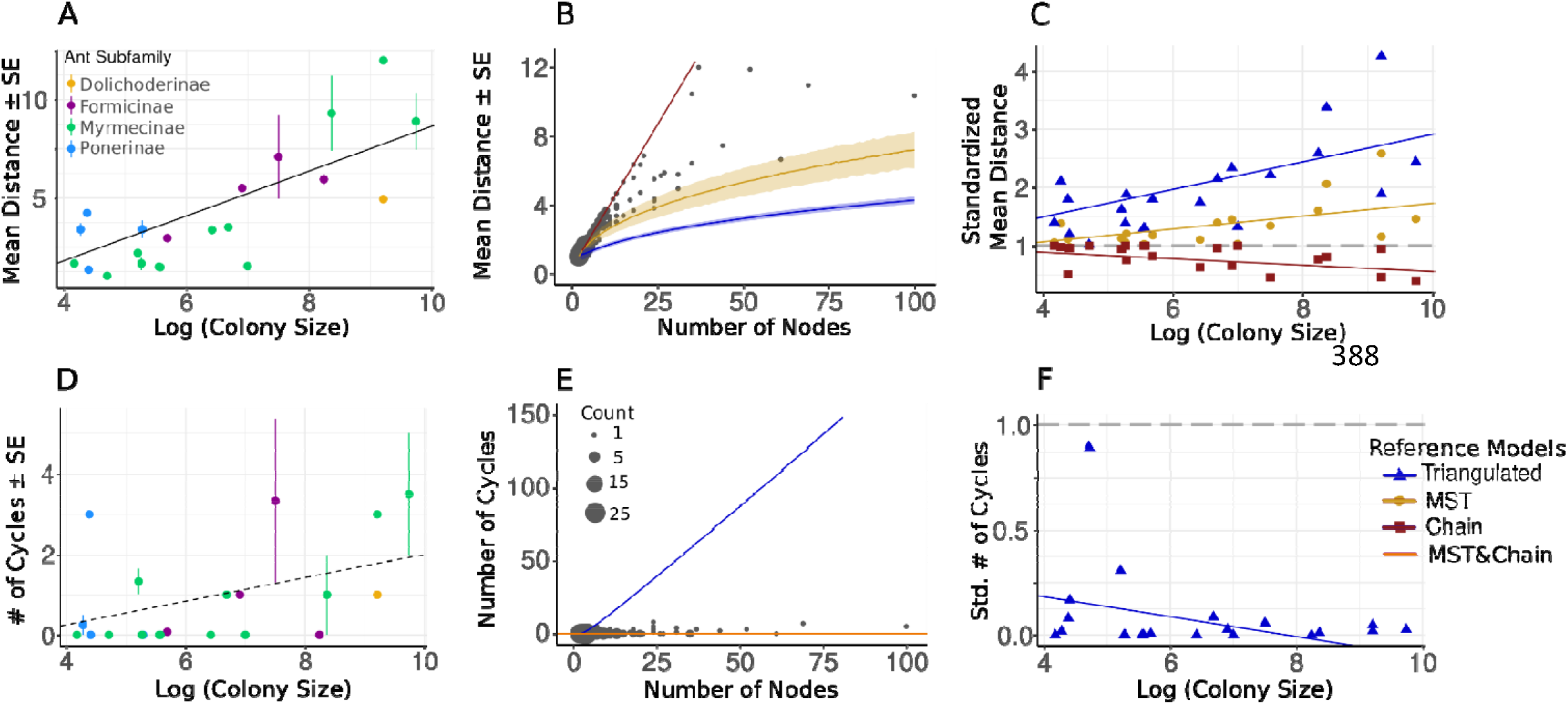
Nest Connectivity: branching (A-C) and short-cut (D-F) hypotheses. (A, D) Relationship between log colony size (average number of ants in a colony) and (A) network mean distance and (D) number of cycles in nests. Each data point represents a mean value for a single species, whiskers indicate standard errors around the mean when multiple nests were sampled for a given species, colors represent ant subfamily (see legend in A). Dashed line is the regression fit for PGLS when λ=1, solid when statistically significant and dashed when not statistically significant; (B, E) The mean distance (B) or number of cycles (E) according to network size (number of nodes). Networks from observed nests are plotted as grey dots, sized according to sample size. In (B) lines are mean distances as a function of number of nodes for chain (red), MST (yellow), and triangulated (blue) reference networks, with 95% confidence intervals from 1000 simulation runs shown as shading around the lines; in (E) Blue line is the number of cycles in triangulated networks as a function of number of nodes. Orange line depicts the number of cycles for both MSTs and chain networks, which contain zero cycles at all network sizes; (C, F) Relationship between log colony size (average number of ants in a colony) and scaled mean distance (C) and scaled number of cycles (F). Mean distance (C) is scaled to chain (red), MST (yellow), and triangulated (blue) networks. Cycles (F) are only scaled to triangulated networks (blue). Each point represents a mean of scaled values from the nests of each species. Grey dashed line at y=1 represents values at which observed nests contain the same value as expected from the reference model. Values below the dashed line indicate networks that have lower values than expected by the reference model. Values above the dashed line indicate networks which have values greater than expected according to the reference model. Solid colored lines indicate statistically significant regression fits for PGLS (λ=1).

### Short-cut Hypothesis

The “short-cut hypothesis,” that nests from species with larger colonies would contain more cycles, was not supported. The relationship between the number of cycles in a nest had a positive relationship with colony size, but this trend was not statistically significant in the majority of the data subsets (Figure 3D; 15/16 subsets; N = 21 genera, OLS of PIC: β =0.293, 95% CI = −0.089, 0.528, R^2^ = 0.160, *p* = 0.072; Table S12). The triangulated reference networks showed a steep increase in the number of cycles as a function of network size, and far exceeded the number of cycles in observed nests given their networks’ size (Figure 3E). However, the number of cycles in observed nests was similar to that expected from the chain and MST reference networks, which do not produce cycles at any network size (Figure 3E). The measure of cycles scaled to the triangulated reference models significantly decreased with colony size in all species subsets (16/16 subsets; Figure 3F; OLS of PIC: β = −0.048, 95% CI = −0.102, −0.015, R^2^ = 0.090, *p* = 0.186; Tables S13).

### Phylogenetic Signal

We detected a significant phylogenetic signal for nest connectivity traits (cycles and mean distance) but not for colony size or nest subdivision traits (chamber number and size). Most traits, except colony size and number of cycles, tended to fit a Brownian Motion model of evolution, however the differences in fit between the alternative models (no signal and estimated Pagel’s lambda) were not significant (<4 AIC units). We did not detect a significant phylogenetic signal for colony size (Table S14; *p =* 0.348, λ = 0.440, AIC_λ_= 99.03) in any species subsets (0/19). Colony size had a slightly better fit to the no signal model (AIC_0_ = 97.29) relative to the Brownian Motion model (AIC_BM_ = 99.35) in all species subsets (19/19 subsets). We did not detect a significant phylogenetic signal for chamber number (Table S15; *p =* 0.091, λ = 1.07, AIC_λ_= 182.72) in all species subsets (19/19). Chamber number had a slightly better fit to the Brownian Motion model (AIC_BM_ = 180.09) relative to the no signal model (AIC_0_ = 182.60) in all species subsets (19/19 subsets). We did not detect a significant phylogenetic signal for standardized chamber size (Table S16; *p =* 0.084, λ = 0.71, AIC_λ_= 88.79) in the majority of species subsets (12/15). Chamber size had a slightly better fit to the Brownian Motion model (AIC_BM_ = 86.83) relative to the no signal model (AIC_0_ = 89.02) in most species subsets (13/15). Nest mean distance normalized to a chain network had a significant phylogenetic signal (Table S17; *p =* 0.001, λ = 1.12, AICλ= −2.22) and had a slightly better fit to the Brownian Motion model (AIC_BM_ = −4.97) relative to the no signal model (AIC_0_ = −2.19) in all species subsets (16/16). The number of cycles had a significant phylogenetic signal (Table S18; *p =* 0.005, λ = 1.12, AIC_λ_=74.97), however we could not distinguish between the fit with the Brownian Motion model (AIC_BM_ = 72.23) and the no signal model (AIC_0_ = 72.39) in most species subsets (13/16).

## DISCUSSION

Our results support the adaptive hypothesis that nest architecture mitigates the challenges of larger colony sizes, i.e., more individuals in a colony. Our comparative analysis found that nests of species with larger colonies are more subdivided and reduce travel distances through increased branching. Overall, we did not find a strong effect of phylogeny, but our findings lay out a foundation for future experimental work on the evolution of mechanisms that allow ant colonies to optimize the structures of their nests.

As predicted by the subdivision hypothesis, the number of chambers was greater in species with larger colonies and chamber size was not related to colony size (Figure 2). Thus, ant nests have evolved from the basal single-chambered form to accommodate larger colonies by adding repeating modules of similar sized chambers rather than increasing chamber (module) size. These results suggest that nests of species with large colonies are divided into more sub-units than nests of species with small colonies. This pattern of increased spatial subdivision in ant nests is shared across a variety of living systems, particularly those that undergo an evolutionary increase in body size. For instance, the mammalian brain subdivides into more cortical units in lineages with larger brain sizes (Kaas 2000, 2017). At smaller biological scales, similar patterns of increased spatial subdivision form the basis for the evolution of multi-cellularity that made larger body sizes possible (Bonner 1998). Even man-made networks, like cities become more subdivided as they increase in size (Anas et al. 1998). Given our correlational findings, it remains unclear if subdivision in nests serves a similar function as it does in these other systems. Further data are needed on the distribution of individuals throughout nest chambers to determine if ants utilize chambers in a way that is consistent with subdivision (e.g., Tschinkel 1999; Murdock and Tschinkel 2015). Nest subdivision may facilitate division of labor through spatial segregation of tasks (Tofts and Franks 1992; Richardson et al. 2011; Mersch et al. 2013), and slow down the spread of disease (Pie et al. 2004; Stroeymeyt et al. 2014, 2018). However, the only direct test of the benefit of nest subdivision showed that subdivision does not increase brood rearing efficiency (Cassill et al. 2002; Tschinkel 2018). Limits on chamber size may also be imposed by physiological processes such as gas exchange or temperature regulation (Jones and Oldroyd 2006), or by structural constraints, because large chambers may collapse. Thus, the causes and consequences of nest subdivision remain to be tested.

The connectivity of nests changed according to colony size, but only one of our hypotheses about nest connectivity was supported. The nests of species with larger colonies had reduced travel distances due to their branching structure, consistent with the branching hypothesis. In the nests of species with small colonies (i.e., up to ~1000 workers in nests containing up to ~10 nodes), mean distances tended to match the values expected for chain networks, and indeed the nests of many small colony species were chains. In the nests of species with large colonies, mean travel distances were shorter than would be expected for chain networks but greater than expected for MSTs, and far greater than expected for triangulated networks (Figure 3B, C). These findings indicate that the nests of species with large colonies have increased branching (i.e., are more tree-like) relative to the nests of species with smaller colonies. These findings show that nest branching evolves in association with colony size, and that increased branching is not necessarily a more “derived” trait or explained by phylogenetic relatedness alone. For example, the nests of some early branching species (e.g. *Dinoponera*) have relatively more branching than the nests of some later branching species (e.g. leaf-cutters, *Mycetagrocious*; Figures 3C & 4). By increasing the frequency of branching, large colonies may curb rising travel distances as their nests increase in size (i.e., contain more chambers). Networks are subject to an inherent tradeoff, in which they cannot simultaneously minimize travel distances (i.e., be highly interconnected) and minimize their number of edges (i.e., minimize infrastructure costs). Increased branching is a common way for biological transport networks to balance this tradeoff (e.g., ant foraging trails: Buhl et al. 2009; slime molds: Tero et al. 2010; neural networks: Raj and Chen 2011; plant growth forms: Conn et al. 2017), especially as body size, or population size, increases (Ringo 1991; Banavar et al. 1999; Ercsey-Ravasz et al. 2013; Cabanes et al. 2015). While these previous studies have identified branching as a way to balance the connectivity-infrastructure cost tradeoff in single species (i.e., over development or between individuals), ours is the first study to show this pattern across lineages in an evolutionary context. Branching is thus a general low-cost way for biological transport networks, including ant nests, to maintain connectivity as they increase in size.

**Figure 4:**
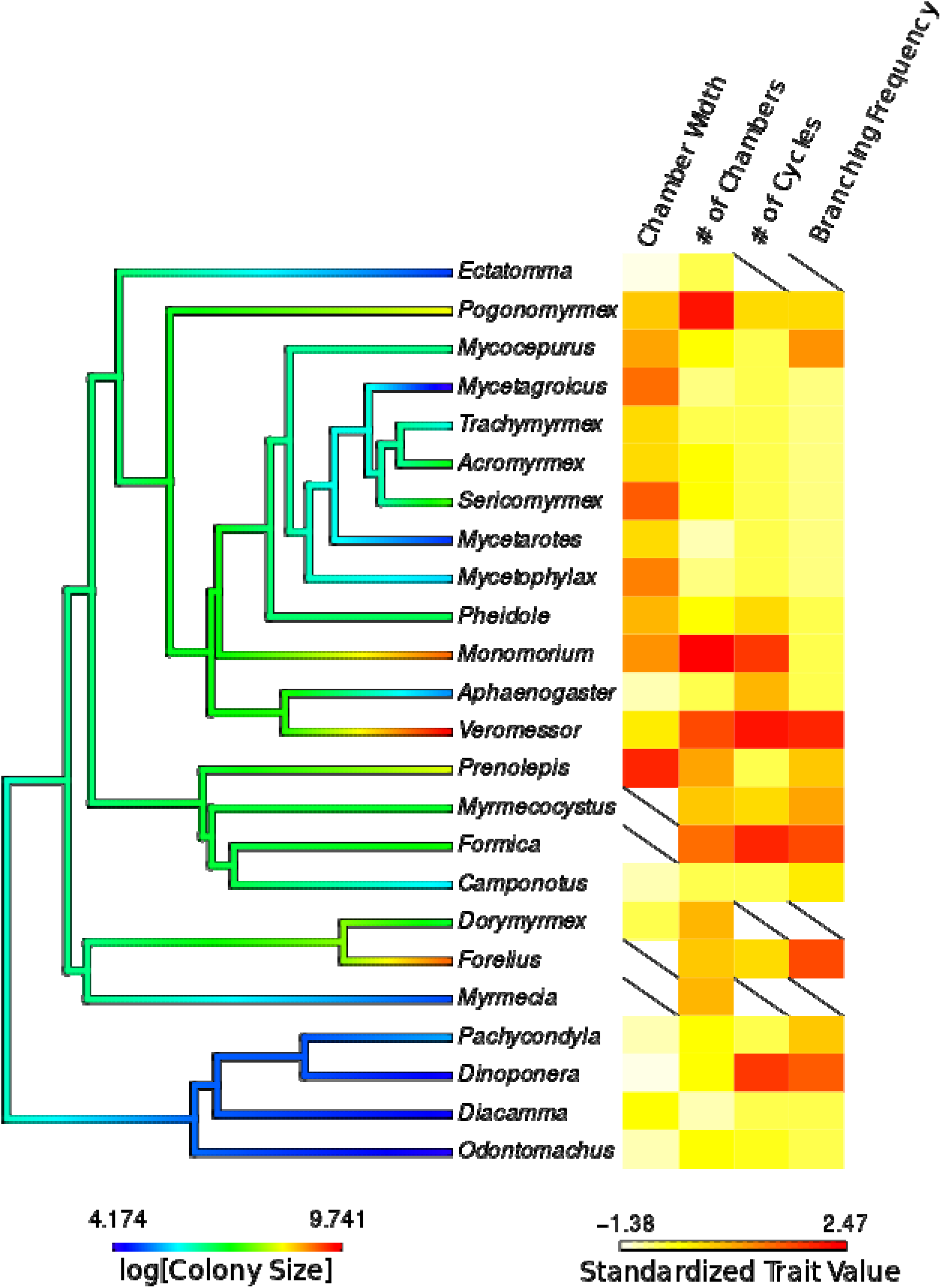
Nest traits and colony size mapped to a genus-level phylogeny of the ants for all genera on which we had data. The log of colony size is mapped onto the branches of the phylogeny, with cooler colors indicating smaller colony sizes and warmer colors larger colony sizes. Values for the four nest traits we analyzed are depicted as a heatmap, with color indicating trait value standardized as a z-score. Branching frequency is a measure of how much nests branch relative to their size; such that values are nest Mean Distance standardized to Chain networks with values multiplied by −1 to ease interpretation, so that larger values indicate greater branching. Data are displayed for species subset one, and slashes indicate missing data.

The short-cuts hypothesis, that nests from species with larger colonies will have more cross-linking paths or cycles, was not supported. Although the number of cycles in a nest increased with colony size, this increase was not statistically significant (Figure 3D). Cycles were added at a slower rate than expected for a triangulated network (Figure 3E), which is not surprising given that triangulations represent the maximum rate at which new cycles could be added with increasing network size (for 2D networks). This finding lends further support to the conclusion that the increase in nest connectivity with colony size is due to increased branching and not the addition of short-cuts. While we did not find support for this adaptive function of cycles, our results lend some support to the hypothesis that variation in cycles is due to phylogenetic processes, like drift, evolutionary history, etc., as cycle number had a significant phylogenetic signal. However, we caution that our analysis conservatively excludes nests that might have the most cycles because we could not accurately infer the topology of the most convoluted nests. Our inability to detect a significant increase in the number of cycles with colony size could be due to this bias in our dataset. More refined imaging techniques could provide a solution to this issue and expand future work on nest architecture in general.

The scarcity of cycles in the nests we measured suggest that the benefits of having alternative access routes, which help maintain movement (Corson 2010; Katifori et al. 2010; Blonder et al. 2018, Jyothi et al. 2016) and facilitate forager recruitment (Pinter-Wollman 2015a), might not outweigh the potential costs of additional tunnels. For instance, there are costs to building and maintaining tunnels (da Silva Camargo et al. 2013; Monaenkova et al. 2015) and there may be negative consequences of many cross-linking paths, including structural instability or heightened exposure to risks, like pathogens (Scovell 1983; Viana et al. 2013; Stroeymeyt et al. 2014, 2018), parasites (Scovell 1983), or predators (Jeanne 1975; LaPolla et al. 2002). Nests are multi-purpose structures that have other functions, like thermoregulation, humidity control or ventilation (Jones and Oldroyd 2006; King et al. 2015), which might suffer when nests have many cycles. In systems where the costs of cross-linking paths are low, like the plasmodia of slime molds, the mycelia of fungus, or the pheromone trails of ants, cycles are ephemeral and created for exploration, yet they are pruned once resources are discovered (Bebber et al. 2007; Buhl et al. 2009; Tero et al. 2010; Latty et al. 2011). Conversely, when the benefits to movement are very high, cycles become more common as size increases, but this only appears to be true in some man-made networks, like urban roads or computer routing networks (Levinson 2012; Jyothi et al. 2016). Future investigation into the building costs of tunnels and the functional consequences of interconnected nest structures will help reveal why ant nests do not contain more short-cuts, or cycles.

While we found support for our adaptive hypotheses, we also found that phylogenetic relatedness explains some of the variation in ant nest structural diversity. Nest traits relating to connectivity (number of cycles and mean distance normalized to chain networks), had significant phylogenetic signals, whereas traits relating to subdivision (chamber number and size) did not. We caution, however, that we did not detect a strong effect of phylogeny overall. The results of our regression analyses did not differ when we assumed a strong phylogenetic signal (λ=1) or none at all (λ=0). Furthermore, the differences in fit between alternative models of evolution (no signal, Brownian Motion, and estimated Pagel’s lambda) were rarely large enough to robustly distinguish between models, which may be due to lack of statistical power. Notably, we did not detect a significant phylogenetic signal for colony size, even though prior studies of ant colony size suggest that phylogenetic relatedness is a factor in colony size evolution (Boulay et al. 2014; Ferguson-gow et al. 2014; Burchill and Moreau 2016). While we cannot reach robust conclusions about the extent of phylogenetic effect on the traits we measured, the co-evolutionary patterns we analyze here account for potential phylogenetic non-independence.

As with any meta-analysis, our dataset is constrained by the available literature. Many of the nests we found in the literature were built in sandy soils, and so our analysis has an over-representation of species adapted to the climate and ecology of sandy soils (Table S1). Fungus growers are highly-represented in our analysis (represented by the crown spanning from *Mycocepurus* to *Mycetophylax* in Figure 4), and constraints imposed by requirements to maintain healthy fungus gardens could have influenced our findings. Additionally, because we do not have data on the size or age of the colonies that built the majority of nests in our meta-analysis, we might have included immature colonies that have not reached their full size. Further work could investigate whether nest connectivity changes throughout development within a species in natural environments (e.g., Tschinkel 1987, 2004, 2005, 2011) or whether there are species-specific connectivity patterns that can be found at different ages.

In conclusion, we show that ant nests exhibit modular structures that minimize travel distancesf in species with larger colonies. The similarity in how ant nests and the bodies of unitary organisms respond to increasing size suggest that their shared architectural features, branching transportation networks and compartmentalization, provide common solutions to the challenges created by large size across the tree of life.

## Acknowledgements

We are thankful to Doug Booher for sharing his nest castings and to Walter Tschinkel for additional images of his published castings. We thank Dana Williams, Felipe Zapata, Kevin Loope, Tyler McCraney and the UCLA IDRE statistical consulting unit for advice on the statistical analysis and to Saray Shai for advice on generating reference models. We thank Gabby Najm, James Lichtenstein, Natalie Lemanski, Andrea Perna, and two anonymous reviewers for helpful comments on the manuscript. The authors declare no conflicts of interest.

