## Supplemental Materials for "Modularity and connectivity of nest structure scale with colony size"

**Figure S1**: (A-E) Images of a 3D scan of a casting from a *Veromessor andrei* nest, shown from different angles to illustrate the different nest features. Labels on each nest feature relate to their position within the network shown in (F). Labels: E – nest entrance; C1-C6 - chambers; J - tunnel junction; END - a tunnel that ends without a chamber. (F) is the network representation of the nest. Edge lengths are not indicative of actual tunnel lengths.

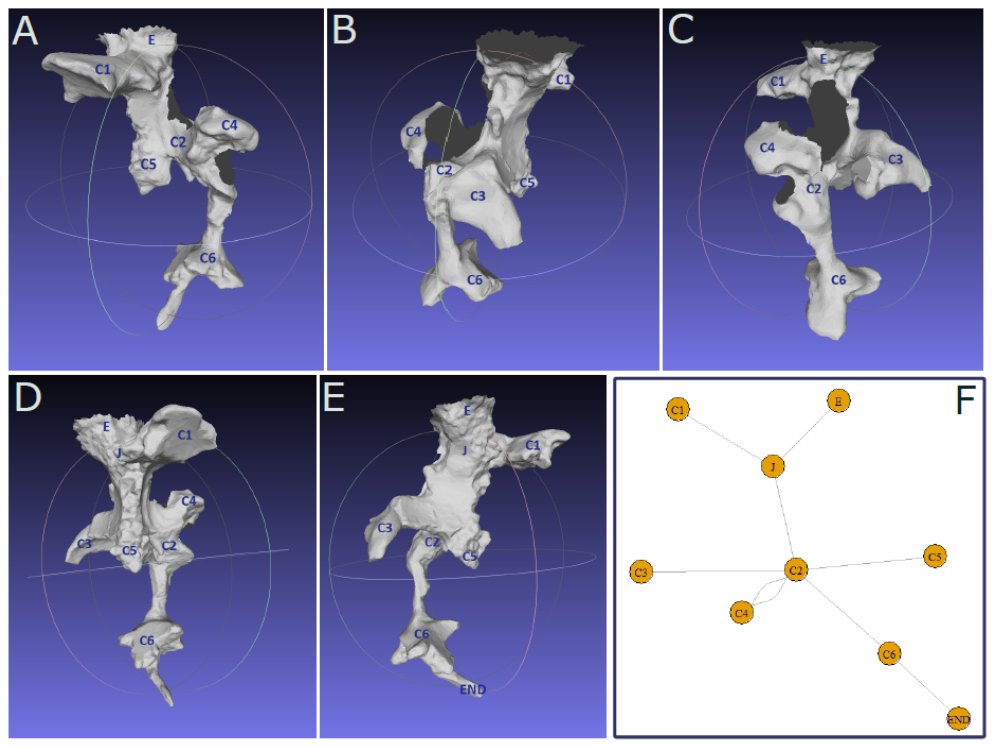

*Chamber Width Validation*

We validated that chamber width was a reliable indicator of chamber area using an independent dataset. We measured the width and area of ten nest chambers from nine castings of *Verromessor andrei* nests from Pinter-Wollman (2015). These castings were not included in our the dataset used for the meta-analysis because they were only partial castings of larger nests. (Pinter-Wollman 2015). We measured one chamber per nest casting, with the exception of one nest casting for which we measured two chambers. A chamber was selected for measurement if at least one of its surfaces (ceiling or floor) was entirely accessible and the plaster completely filled the floor of the chamber, as indicated by soil markings on the vertical edges (chamber walls) of the chamber casting. To then obtain a precise measure of chamber area, we created a mold of the chamber floor by covering the surface in modeling clay. We then extracted the clay mold and photographed it with a ruler for scale. We measured area and maximum width of the mold from photographs using ImageJ (Rueden et al. 2017). To validate that our measure of maximum width is a reliable indicator of chamber area, we compared the two measures - width and area - using a nonlinear regression with quadratic fit (y= β_1_x+ β_2_x^2^+α). Chamber width was a strong and significant predictor of chamber area (β_1_ = 1.34 ± 3.89, β_2_ = 0.40 ± 0.24, R^2^ = 0.97, p <0.001) for the range of chamber widths measured (width mean = 7.28 cm, SD = 2.75 cm, range = 4.0 -12.0 cm). The widths of nest chambers in our dataset are well represented by the range of the validation subset (mean = 7.32 cm, SD = 4.33 cm, range = 1.0-38.0 cm).

**Table S1**: Species used for each analysis, including information on the number of nests measured per species,

average colony size, and a note about each species’ natural history. Source references for nest images and colony

size values are provided in the last two columns.

| **Species Name** | **# Nests** | **Used in Connect-ivity Analysis** | | **Used in Sub-**  **division Analysis**  **Chamber #** | | **Used in Sub-**  **division Analysis**  **Chamber Size** | | **Colony Size (Avg)** | | **Workers**  **Monomorphic or Polymorphic** | | **Natural History Notes** | | **Nest Structure Source** | | **Colony Size Source** |
| --- | --- | --- | --- | --- | --- | --- | --- | --- | --- | --- | --- | --- | --- | --- | --- | --- |
| *Acromyrmex balzani* | 15 | Y (14) | Y | | Y | | 1095 | | Polymorphic | | Grass-cutting fungus grower found in cerrado of South America. | | Caldato, N., da et al. 2016. J. of Nat. Hist., 50(25–26): 1561–1581. | | Caldato, N., de Andrade, A. P., Forti, L. C., Barbieri, R. F. & Lopes, J. F. S. 2010. Sociobiol., 56(3): 727–736. | |
| *Acromyrmex landolti* | 7 | N | Y | | Y | | 1000 | | Polymorphic | | Leaf-cutting fungus grower found in semi-arid South American Caatinga. | | Verza, S. S., et al. 2020. Insectes Sociaux 67(1): 147–153. | | Beckers, R., Goss, S., Deneubourg, J., & Pasteels, J.1989. Psyche, 96: 239–256. | |
| *Aphaenogaster treatae* | 11 | Y | Y | | Y | | 168 | | Monomorphic | | Forages on arthropods and seeds in long leaf pine savanna. | | Tschinkel, W. R. 2011. J. Insect Science 11:1–30. | | Tschinkel, W. R. 2011. J. Insect Science 11:1–30 | |
| *Aphaenogaster floridana* | 21 | Y | Y | | Y | | 183 | | Monomorphic | | Forages on arthropods and seeds in sandy soils including long leaf pine savanna. | | Tschinkel, W. R. 2011 J. Insect Science 11:1–30. | | Tschinkel, W. R. 2011. J. Insect Science 11:1–30. | |
| *Aphaenogaster ashmeadi* | 11 | Y | Y | | Y | | 137 | | Monomorphic | | Forages on arthropods in sandy soils in pine, oak and oak scrub habitats. | | Tschinkel, W. R. 2011. J. Insect Science 11:1–30 | | Tschinkel, W. R. 2011. J. Insect Science 11:1–30 | |
| *Camponotus socius* | 13 | Y | Y | | Y | | 296 | | Polymorphic: continuous | | Generalist foragers, mass recruitment, in sandy soils, including long leaf pine savanna. | | Tschinkel, W. R. 2005. J. Insect Science 5:1–18. | | Tschinkel, W. R. 2005. J. Insect Sci. 5:1–18. | |
| *Diacamma indicum* | 13 | Y | Y | | Y | | 82 | | Monomorphic | | Solitary termite hunters and scavengers from India, exposed to seasonal flooding. | | Bhattacharyya, K., & Annagiri, S. 2019. Journal of Insect Science, 19(6). | | Bhattacharyya, K., & Annagiri, S. 2019. Journal of Insect Science, 19(6). | |
| *Dorymyrmex bureni* | 1 | N | Y | | Y | | 1,000 | | Monomorphic | | Group and solitary foragers, on live arthropods and nectar, sandy soils in Southeastern US | | Tschinkel, W. R. 2003. Palaeogeogr. Palaeoclimatol. Palaeoecol. 192:321–333. | | King, J. R. 2010. Ecol. Entomol. 35:287–298. | |
| *Dinoponera australis* | 1 | Y | Y | | N | | 14 | | Monomorphic | | Solitary hunters of arthropods in tropical South America. | | Paiva, R. V. S., & Brandão, C. R. F. 1995. Ethology Ecology and Evolution, 7(4), 297–312. | | Paiva, R. V. S., & Brandão, C. R. F. 1995. Ethology Ecology and Evolution, 7(4), 297–312. | |
| *Dinoponera quadriceps* | 1 | Y | Y | | Y | | 80 | | Monomorphic | | Solitary scavengers and hunters of arthropods in tropical savanna, moist and wet forests in Atlantic Forest | | Vasconellos, A., et al. 2004. Brazilian J. Biol. 64:357–362. | | Vasconellos, A., G. G. Santana, and A. K. Souza. 2004. Brazilian J. Biol. 64:357–362. | |
| *Ectatomma brunneum* | 5 | N | Y | | Y | | 116 | | Monomorphic | | Solitary hunters of arthropods, broad South American distribution | | Renard, D., et al. 2010. Ecoscience, 17, (2), 194–202. | | Vieira, A. S., and W. F. Antonialli Junior. 2006. Sociobiology 47:275–287. | |
| *Forelius sp.* | 1 | Y | Y | | N | | 10,000 | | Monomorphic | | Solitary and group foragers on live and dead arthropods and nectar in Southwestern US | | Doug Booher's casting | | Kaspari, M., and T. J. Valone. 2002. Ecology 83:2991–2996. | |
| *Formica archboldi* | 3 | Y | Y | | Y(1) | | 500 | | Monomorphic | | Forages on live arthropods and tends nectar in upland pine and scrub of Southeastern US. | | King, J. R., and J. C. Trager. 2007. J. Insect Sci. 7:1–14.; Tschinkel, W. R. 2015. J. Bioeconomics 17:271–291 | | King, J. R., and J. C. Trager. 2007. J. Insect Sci. 7:1–14. | |
| *Formica dolosa* | 1 | Y | Y | | Y | | 600 | | Monomorphic | | Generalist scavengers, host to dulotic *Polyergus longicornis*, in Eastern US. | | King, J. R., and J. C. Trager. 2007. J. Insect Sci. 7:1–14. | | King, J. R., and J. C. Trager. 2007. J. Insect Sci. 7:1–14. | |
| *Formica pallidefulva* | 1 | Y | Y | | Y | | 400 | | Monomorphic | | Generalist scavengers, host to several dulotic social parasites, broad distribution in North America. | | Tschinkel, W. R. 2003. Palaeogeogr. Palaeoclimatol. Palaeoecol. 192:321–333. | | King, J. R., and S. D. Porter. 2007. Evol. Ecol. Res. 9:757–774. | |
| *Formica japonica* | 4 | Y | Y | | N | | 1811 | | Monomorphic | | Generalist scavengers, recruit to foraging trails. Broad distribution in East Asia | | Kondoh, M. 1968. Ecol. Soc. Japan 18:124–133. | | Kondoh, M. 1968. Ecol. Soc. Japan 18:124–133. | |
| *Formica subaenescens* | 1 | Y | Y | | Y | | 1246 | | Monomorphic | | Generalist foragers, broad distribution in North America. | | Markin, G. P. 1964. Ann. Entomol. Soc. Am. 57:360–362. | | Tuzzolino, K. E., and W. D. Brown. 2010. Entomol. Sci. 13:162–165. | |
| *Monomorium viridum* | 1 | Y | Y | | Y | | 10,000 | | Monomorphic | | Generalist foragers, broad distribution in North America. | | Williams, D. F. and C. S. Lofgren. 1988. Pp. 433–443 *in* Advances in Myrmecolog. | | King, J. R., and S. D. Porter. 2007. Evol. Ecol. Res. 9:757–774. | |
| *Mycetagroicus inflatus* | 4 | Y | Y | | Y | | 65 | | Monomorphic | | Fungus-grower in alluvial forests in Para, Brazil | | Ješovnik, A., et al. 2013. Insectes Soc. 60:531–542. | | Ješovnik, et al. 2013. Insectes Soc. 60:531–542. | |
| *Mycetarotes acutus* | 5 | Y | Y | | Y | | 25 | | Monomorphic | | Fungus-grower on seeds and plant fibers in Brazil and Ecuador | | Solomon, S. E., et al. *et al.* 2004. Insectes Soc. 51:333–338. | | Mayhé-nunes, A. J., and C. R. Ferreira Brandão. 2006. Rev. Bras. Entomol. 50:463–472. | |
| *Mycetarotes parallelus* | 19 | Y | Y | | Y | | 111 | | Monomorphic | | Fungus-grower in South America | | Solomon, S. E., *et al.* 2004. Insectes Soc. 51:333–338. | | Solomon, S. E., *et al.* 2004. Insectes Soc. 51:333–338. | |
| *Mycetophylax simplex* | 8 | Y | Y | | Y(6) | | 264 | | Monomorphic | | Fungus-grower along sandy beaches of the Atlantic Forest in Brazil and Uruguay | | Klingenberg, C., et al. 2007. Stud. Neotrop. Fauna Environ. 42:121–126.; Diehl-fleig, E., E. Diehl. 2007. Insectes Soc. 54:242–247. | | Diehl-fleig, E., and E. Diehl. 2007. Insectes Soc. 54:242–247. | |
| *Mycocepurus goeldii* | 1 | Y | Y | | N | | 613 | | Monomorphic | | Fungus-grower on caterpillar frass, broad distribution in South America | | Rabeling, C., M. et al. 2007. J. Insect Sci. 7:1–13. | | Rabeling, C., M. et al. 2007. J. Insect Sci. 7:1–13. | |
| *Mycocepurus smithii* | 1 | Y | Y | | Y | | 77 | | Monomorphic | | Fungus-grower on frass and dead plant matter, broad distribution in Central and South America | | Rabeling, C., M. et al. 2007. J. Insect Sci. 7:1–13.. | | Marín, H. F. J. K. Zimmerman, W. T. Wcislo, S. A. Rehner. 2005. J. Nat. Hist. 39:1735–1743. | |
| *Myrmecia dispar* | 1 | Y | Y | | N | | 124 | | Monomorphic | | Solitary foragers on live arthropods and nectar in inland forests and heath of Southern Australia. | | Gray, B. 1971. Insectes Sociaux, 18: 71–80. | | Gray, B. 1971. Insectes Sociaux, 18: 71–80. | |
| *Myrmecocystus navajo* | 1 | Y | Y | | N | | 1000 | | Polymorphic: continuous | | Solitary foragers on nectar and dead arthropods in deserts of the Southwestern US. | | Doug Booher's casting | | [http://www.antwiki.org/wiki/Myrmecocystus_navajo, accessed August 2018](http://www.antwiki.org/wiki/Myrmecocystus_navajo,%20accessed%20August%202018) | |
| *Odontomachus brunneus* | 12 | Y | Y | | Y | | 72 | | Monomorphic | | Solitary hunters of arthropods in the Southeastern US | | Cerquera, L. M., and W. R. Tschinkel. 2008. J. Insect Sci. 10:1–12.; Tschinkel, W. R. 2003. Palaeogeogr. Palaeoclimatol. Palaeoecol. 192:321–333. | | Cerquera, L. M., and W. R. Tschinkel. 2008. J. Insect Sci. 10:1–12. | |
| *Odontomachus chelifer* | 5 | Y(1) | Y | | N | | 499 | | Monomorphic | | Solitary hunters of arthropods, broad distribution in Central and South America | | Guimarães, I. de C., et al. 2018. PLoS One 13:e0189896. | | Guimarães, I. de C., et al. 2018. PLoS One 13:e0189896. | |
| *Pachycondyla striata* | 4 | Y | Y | | Y | | 197 | | Monomorphic | | Solitary foragers on arthropods and seed elaiosomes, in southern Atlantic rainforests and cerrado of South America. | | da Silva-Melo, A., & Giannotti, E. 2010. Insectes Sociaux, 57: 17–22. | | da Silva-Melo, A., & Giannotti, E. 2010. Insectes Sociaux, 57: 17–22. | |
| *Pheidole dentata* | 1 | Y | Y | | Y | | 800 | | Polymorphic: discrete | | Southeastern US | | Williams, D. F. and C. S. Lofgren. 1988. Pp. 433–443 *in* Advances in Myrmecolog. | | King, J. R., and S. D. Porter. 2007. Evol. Ecol. Res. 9:757–774. | |
| *Pogonomyrmex badius* | 4 | Y(3) | Y | | Y | | 4,300 | | Monomorphic | | Seed harvester, recruits to foraging trails in sandy soils of Southeastern US | | Tschinkel, W. R. 2014. PLoS One 9 (11): e112981. <https://youtu.be/7XngDAiOQC4?list=PLoUFkrkM_ZDcLqVV4TGjDlfplFmWsFJI7> and <https://youtu.be/nAdZSyTYsmU?list=PLoUFkrkM_ZDcLqVV4TGjDlfplFmWsFJI7> and <https://youtu.be/fcbYezl02c4?list=PLoUFkrkM_ZDcLqVV4TGjDlfplFmWsFJI7> | | Tschinkel, W. R. 2004. J. Insect Sci. 21:1–19. | |
| *Prenolepis imparis* | 1 | N | Y | | Y | | 3,790 | | Monomorphic | | Generalist on arthropods and nectar, recruit to foraging trails, broad distribution in North America | | Tschinkel, W. R. 1987. Insectes Soc. 34:143–164. | | Tschinkel, W. R. 1987. Insectes Soc. 34:143–164. | |
| *Sericomyrmex amabilis* | 2 | Y(1) | Y | | Y | | 471 | | Polymorphic | | Fungus-grower on organic debris in Central America and northern South America. Host to ant social parasite, *Megalomyrmex symmetochus.* | | Ješovnik, A., et al. 2018. Myrmecol. News: 26, 65–80. | | Ješovnik, A., et al. 2018. Myrmecol. News: 26, 65–80. | |
| *Sericomyrmex bondari* | 2 | Y | Y | | Y | | 845 | | Polymorphic | | Fungus-grower on organic debris, broad distribution in northern South America. | | Ješovnik, A., et al. 2018. Myrmecol. News: 26, 65–80. | | Ješovnik, A., et al. 2018. Myrmecol. News: 26, 65–80. | |
| *Seriocomyrmex mayri* | 5 | Y(1) | Y | | Y | | 1580 | | Polymorphic | | Fungus-grower on organic debris, broad distribution in northern South America. | | Ješovnik, A., et al. 2018. Myrmecol. News: 26, 65–80. | | Ješovnik, A., et al. 2018. Myrmecol. News: 26, 65–80. | |
| *Seriocomyrmex opacus* | 2 | N | Y | | Y | | 168 | | Polymorphic | | Fungus-grower on organic debris in forests of Central America and northern South America. | | Ješovnik, A., et al. 2018. Myrmecol. News: 26, 65–80. | | Ješovnik, A., et al. 2018. Myrmecol. News: 26, 65–80. | |
| *Seriocomyrmex parvulus* | 3 | Y(2) | Y | | Y | | 258 | | Polymorphic | | Fungus-grower on organic debris, broad distribution in forests of northern South America. | | Ješovnik, A., et al. 2018. Myrmecol. News: 26, 65–80. | | Ješovnik, A., et al. 2018. Myrmecol. News: 26, 65–80. | |
| *Seriocomyrmex saramama* | 1 | Y | Y | | Y | | 51 | | Polymorphic | | Fungus-grower on organic debris in western South America. | | Ješovnik, A., et al. 2018. Myrmecol. News: 26, 65–80. | | Ješovnik, A., et al. 2018. Myrmecol. News: 26, 65–80. | |
| *Sericomyrmex saussurei* | 1 | N | Y | | Y | | 1249 | | Polymorphic | | Fungus-grower on organic debris, broad distribution in forests and cerrado of northern South America. | | Ješovnik, A., et al. 2018. Myrmecol. News: 26, 65–80. | | Ješovnik, A., et al. 2018. Myrmecol. News: 26, 65–80. | |
| *Trachymyrmex holmgreni* | 50 | Y(1) | Y | | Y(1) | | 383 | | Polymorphic | | Fungus-grower on grass, nests in sandy soils, broad South American distribution. | | Cristiano M.P., et al. 2019 Insectes Soc. 66, 139–151. | | Cristiano MP, et al. 2019 Insectes Soc. 66, 139–151. | |
| *Trachymyrmex septentrionalis* | 2 | Y | Y | | Y | | 194 | | Monomorphic | | Fungus-grower on frass and dead plant matter in sandy soils of Eastern US | | Tschinkel, W. R. 2015. J. Bioeconomics 17:271–291; and  Tschinkel, W. R. 2003. Palaeogeogr. Palaeoclimatol. Palaeoecol. 192:321–333. | | Seal, J. N., W. R. Tschinkel. 2006. Ann. Entomol. Soc. Am. 99:673–682. | |
| *Veromessor pergandei* | 1 | Y | Y | | Y | | 17,000 | | Polymorphic: continuous | | Desert seed harvester in Southwestern US. | | Doug Booher's casting | | Went, F. W., J. Wheeler, and G. C. Wheeler. 1972. Bio-science 22:82-88 | |

**Table S2**: Excluded Papers

| **Reference** | **Reason Excluded** |
| --- | --- |
| Antonialli, W. F. and E. Giannotti. 1997. Nest architecture and population dynamics of the ponerine ant Ectatomma edentatum (Hymenoptera, Formicidae). J. ofAdvanced Zool. 18:64–71. | Polydomous and sets of nests not identified to single colonies |
| Cole, B. J. 1994. Nest architecture in the western harvester ant , Pogonomyrmex occidentalis (Cresson ). Insectes Sociaux 41:401–410. | Missing useful images or tables |
| Conway, J. R. 1983. Nest Architecture and Population of the Honey Ant , Myrmecocystus mexicanus Wesmael ( Formicidae ), in Colorado. Southwestern Association of Naturalists 28:21–31. | Missing useful images or tables |
| Forti, Luiz C., et al. “The nest architecture of the ant, Pheidole oxyops Forel, 1908 (Hymenoptera: Formicidae).” Insect Science, vol. 14, no. 5, 2007, pp. 437–442 | No colony size info for *Pheidole oxyops* |
| Halley JD, Burd M, Wells P. 2005. Excavation and architecture of Argentine ant nests. Insectes Soc. 52:350–356. | Too convoluted |
| Lavigne, R. J. 1969. Bionomics and Nest Structure of Pogonomyrmex occidentalis (Hymenoptera: Formicidae). Annals of the Entomological Society of America 62:1166–1175. | Missing nest tops |
| Lopes, J. F. S., L. F. Ribeiro, M. S. Brugger, R. D. S. Camargo, N. Caldato, and L. C. Forti. 2011. Internal architecture and population size of Acromyrmex subterraneus molestans (Hymenoptera, Formicidae) nests: Comparison between a rural and an urban area. Sociobiology 58:593–605. | No colony size info for *Acromyrmex subterraneus* |
| Minter, N. J., N. R. Franks, and K. A. R. Brown. 2012. Morphogenesis of an extended phenotype :four-dimensional ant nest architecture. Jounral of the Royal Society Inteface 9:586–595. | Artificial excavation conditions |
| Moser, J. C. 2006. Complete Excavation and Mapping of a Texas Leafcutting Ant Nest. Annales of the Entomological Society of America 99:891–897. | Cannot view nest |
| Peeters, C., et al. “"Wall-Papering" and elaborate nest architecture in the ponerine antHarpegnathos saltator.” Insectes Sociaux, vol. 41, no. 2, 1994, pp. 211–218. | Cannot decipher images |
| Scherba, G. 1961. Nest Structure and Reproduction in the Mound-Building Ant Formica opaciventris Emery in Wyoming. New York Entom. Journal of the New York Entolomological Society 69:71–87. | No images or tables with relevant info |
| Talbot, M., and C. H. Kennedy. 1940. The slave-making ant, Formica sanguinea subintegra Emery, its raids, nuptual flights and nest structure. Annals of the Entomological Society of America 33:560–577. | Images are incomplete |
| Tschinkel, W. R. (2004). The nest architecture of the Florida harvester ant, Pogonomyrmex badius. Journal of Insect Science, 4, 21. | Too convoluted |
| Whitford, W. G., P. Johnson, and J. Ramirez. 1976. Comparative ecology of the harvester ants Pogonomyrmex Barbatus (F. Smith) and Pogonomyrmex Rugosus (Emery). Insectes Soc. 23:117–131. | Nest tunnels and interior obscured |

**RESULTS FROM DIFFERENT DATA SUBSETS**

**Nest Subdivision**

Chamber Number

Nine genera had two or more species each represented in our data set for the chamber number analysis: *Acromyrmex, Aphaenogaster,* *Dinopera*, *Formica,* *Mycocepurus, Mycetarotes, Odontomachus,* *Sericomyrmex* and *Trachymyrmex*. Because our analysis was restricted to the genus level, we used a different subset of the species that represent these nine genera in our analysis and report in the main text the proportion of these subsets that were significant. Data subsets with significant results are in bold in the tables below. The list of species subsets contained:

Subset 1: *Acromyrmex balzani, Aphaenogaster floridana, Dinoponera quadriceps, Formica japonica, Mycetarotes parallelus, Mycocepurus goeldii, Odontomachus brunneus, Sericomyrmex mayri, Trachymyrmex holmgreni*

Subset 2: ***Acromyrmex landolti****, A. floridana, D. quadriceps, F. japonica, M. parallelus, M. goeldii, O. brunneus, S. mayri, T. holmgreni*

Subset 3: *A. balzani,* ***Aphaenogaster treatae****, D. quadriceps, F. japonica, M. parallelus, M. goeldii, O. brunneus, S. mayri, T. holmgreni*

Subset 4: *A. balzani,* ***Aphaenogaster ashmeadi****, D. quadriceps, F. japonica, M. parallelus, M. goeldii, O. brunneus, S. mayri, T. holmgreni*

Subset 5: *A. balzani, A. floridana,* ***Dinoponera australis****, F. japonica, M. parallelus, M. goeldii, O. brunneus, S. mayri, T. holmgreni*

Subset 6: *A. balzani, A. floridana, D. quadriceps,* ***Formica archboldi****, M. parallelus, M. goeldii, O. brunneus, S. mayri, T. holmgreni*

Subset 7: *A. balzani, A. floridana, D. quadriceps,* ***Formica dolosa****, M. parallelus, M. goeldii, O. brunneus, S. mayri, T. holmgreni*

Subset 8: *A. balzani, A. floridana, D. quadriceps,* ***Formica pallidefulva****, M. parallelus, M. goeldii, O. brunneus, S. mayri, T. holmgreni*

Subset 9: *A. balzani, A. floridana, D. quadriceps,* ***Formica subaenescens****, M. parallelus, M. goeldii, O. brunneus, S. mayri, T. holmgreni*

Subset 10: *A. balzani, A. floridana, D. quadriceps, Formica japonica,* ***Mycetarotes acutus****, M. goeldii, O. brunneus, S. mayri, T. holmgreni*

Subset 11: *A. balzani, A. floridana, D. quadriceps, F., japonica, M. parallelus,* ***Mycocepurus smithii****, O. brunneus, S. mayri, T. holmgreni*

Subset 12: *A. balzani, A. floridana, D. quadriceps, F. japonica, M. parallelus, M. goeldii,* ***Odontomachus chelifer****, S. mayri, T. holmgreni*

Subset 13: *A. balzani, A. floridana, D. quadriceps, Formica japonica, M. parallelus, M. goeldii, O. brunneus,* ***Sericomyrmex amabilis****, T. holmgreni*

Subset 14: *A. balzani, A. floridana, D. quadriceps, F. japonica, M. parallelus, M. goeldii, O. brunneus,* ***Sericomyrmex bondari****, T. holmgreni*

Subset 15: *A. balzani, A. floridana, D. quadriceps, F. japonica, M. parallelus, M. goeldii, O. brunneus,* ***Sericomyrmex opacus****, T. holmgreni*

Subset 16: *A. balzani, A. floridana, D. quadriceps, F. japonica, M. parallelus, M. goeldii, O. brunneus,* ***Sericomyrmex parvulus****, T. holmgreni*

Subset 17: *A. balzani, A. floridana, D. quadriceps, F. japonica, M. parallelus, M. goeldii, O. brunneus,* ***Sericomyrmex saramama****, T. holmgreni*

Subset 18: *A. balzani, A. floridana, D. quadriceps, F. japonica, M. parallelus, M. goeldii, O. brunneus,* ***Sericomyrmex saussurei****, T. holmgreni*

Subset 19: *A. balzani, A. floridana, D. quadriceps, F. japonica, M. parallelus, M. goeldii, O. brunneus, S. mayri,* ***Trachymyrmex septentrionalis***

**Table S3:** Chamber Number ~ Log(Colony Size), OLS of PIC (N = 24 genera)

| **Data Subset** | **Coefficient (β)** | **95% CI for β (0.025,0.975)** | **R^2^** | **p-value** | **AIC** |
| --- | --- | --- | --- | --- | --- |
| 1 | 4.024 | **2.014, 6.292** | 0.560 | <0.001 | 51.547 |
| 2 | 3.977 | **2.107, 6.108** | 0.551 | <0.001 | 51.672 |
| 3 | 4.074 | **2.162, 6.294** | 0.565 | <0.001 | 51.814 |
| 4 | 4.137 | **1.978, 6.345** | 0.574 | <0.001 | 52.120 |
| 5 | 3.403 | **1.197, 5.520** | 0.422 | <0.001 | 58.609 |
| 6 | 3.917 | **1.925, 6.051** | 0.562 | <0.001 | 49.963 |
| 7 | 3.916 | **1.927, 6.018** | 0.557 | <0.001 | 50.352 |
| 8 | 3.879 | **1.867, 5.956** | 0.549 | <0.001 | 50.762 |
| 9 | 3.974 | **1.890, 6.069** | 0.559 | <0.001 | 50.909 |
| 10 | 3.678 | **1.855, 5.769** | 0.542 | <0.001 | 52.474 |
| 11 | 3.853 | **1.742, 5.821** | 0.547 | <0.001 | 52.304 |
| 12 | 4.389 | **2.251, 7.178** | 0.459 | <0.001 | 64.918 |
| 13 | 4.198 | **1.959, 6.429** | 0.545 | <0.001 | 52.707 |
| 14 | 4.001 | **1.680, 6.307** | 0.516 | <0.001 | 53.881 |
| 15 | 4.342 | **2.058, 6.091** | 0.587 | <0.001 | 50.594 |
| 16 | 4.302 | **2.034, 6.225** | 0.580 | <0.001 | 50.479 |
| 17 | 4.064 | **2.126, 5.941** | 0.580 | <0.001 | 50.986 |
| 18 | 3.941 | **1.819, 6.310** | 0.527 | <0.001 | 53.077 |
| 19 | 4.003 | **2.087, 5.887** | 0.573 | <0.001 | 51.416 |

Chamber Width

Six genera had two or more species each represented in our data set for the chamber size analysis: *Acromyrmex, Aphaenogaster,* *Formica,* *Mycetarotes,* *Sericomyrmex* and *Trachymyrmex*. Because our analysis was restricted to the genus level, we used a different subset of the species that represent these six genera in our analysis and report in the main text the proportion of these subsets that were significant. Data subsets with significant results are in bold in the tables below. The list of species subsets contained:

Subset 1: *Acromyrmex balzani, Aphaenogaster floridana, Formica archboldi, Mycetarotes parallelus, Sericomyrmex mayri, Trachymyrmex holmgreni*

Subset 2: ***Acromyrmex landolti****, A. floridana, F. archboldi, M. parallelus, S. mayri, T. holmgreni*

Subset 3: *A. balzani,* ***Aphaenogaster treatae****, F. archboldi, M. parallelus, S. mayri, T. holmgreni*

Subset 4: *A. balzani,* ***Aphaenogaster ashmeadi****, F. archboldi, M. parallelus, S. mayri, T. holmgreni*

Subset 5: *A. balzani, A. floridana,* ***Formica pallidefulva****, M. parallelus, S. mayri, T. holmgreni*

Subset 6: *A. balzani, A. floridana,* ***Formica dolosa****, M. parallelus, S. mayri, T. holmgreni*

Subset 7: *A. balzani, A. floridana,* ***Formica subaenescens****, M. parallelus, S. mayri, T. holmgreni*

Subset 8: *A. balzani, A. floridana, F. archboldi,* ***Mycetarotes acutus****, S. mayri, T. holmgreni*

Subset 9: *A. balzani, A. floridana, F. archboldi, M. parallelus,* ***Sericomyrmex amabilis****, T. holmgreni*

Subset 10: *A. balzani, A. floridana, F. archboldi, M. parallelus,* ***Sericomyrmex bondari****, T. holmgreni*

Subset 11: *A. balzani, A. floridana, F. archboldi, M. parallelus,* ***Sericomyrmex opacus****, T. holmgreni*

Subset 12: *A. balzani, A. floridana, F. archboldi, M. parallelus,* ***Sericomyrmex parvulus****, T. holmgreni*

Subset 13: *A. balzani, A. floridana, F. archboldi, M. parallelus,* ***Sericomyrmex saramama****, T. holmgreni*

Subset 14: *A. balzani, A. floridana, F. archboldi, M. parallelus,* ***Sericomyrmex saussurei****, T. holmgreni*

Subset 15: *A. balzani, A. floridana, F. archboldi, M. parallelus, S. mayri,* ***Trachymyrmex septentrionalis***

**Table S4:** Standardized Chamber Width~ Log(Colony Size), OLS of PIC (N = 21 genera)

| **Data Subset** | **Coefficient (β)** | **95% CI for β (0.025,0.975)** | **R^2^** | **p-value** | **AIC** |
| --- | --- | --- | --- | --- | --- |
| 1 | 0.325 | -0.086, 0.932 | 0.102 | 0.158 | -4.719 |
| 2 | 0.442 | 0.088, 0.975 | 0.193 | 0.046 | -7.541 |
| 3 | 0.313 | -0.097, 1.009 | 0.097 | 0.170 | -4.944 |
| 4 | 0.315 | -0.073, 0.874 | 0.099 | 0.164 | -4.952 |
| 5 | 0.329 | -0.061, 0.961 | 0.108 | 0.147 | -5.412 |
| 6 | 0.320 | -0.044, 0.910 | 0.098 | 0.168 | -4.437 |
| 7 | 0.295 | -0.086, 0.893 | 0.085 | 0.200 | -4.539 |
| 8 | 0.325 | -0.034, 0.829 | 0.115 | 0.132 | -4.877 |
| 9 | 0.241 | -0.213, 0.822 | 0.055 | 0.307 | -5.881 |
| 10 | 0.321 | -0.071, 0.893 | 0.087 | 0.194 | -3.319 |
| 11 | 0.217 | -0.221, 0.748 | 0.046 | 0.353 | -6.188 |
| 12 | 0.224 | -0.256, 0.704 | 0.042 | 0.376 | -3.309 |
| 13 | -0.053 | -0.854, 0.706 | 0.002 | 0.852 | 4.547 |
| 14 | 0.588 | **0.137, 1.429** | 0.145 | 0.088 | 10.286 |
| 15 | 0.311 | -0.095, 0.983 | 0.089 | 0.189 | -4.675 |

**Nest Connectivity**

Eight genera had two or more species each represented in our data set for the network analyses: *Aphaenogaster,* *Dinoponera, Formica, Mycetarotes, Mycocepurus, Odontomachus, Sericomyrmex* and *Trachymyrmex*. Because our analysis was restricted to the genus level, we used a different subset of the species that represent these eight genera in our analysis and report in the main text the proportion of these subsets that were significant. Data subsets with significant results are in bold in the tables below. The list of species subsets contained:

Subset 1: *Aphaenogaster floridana, Dinoponera quadriceps, Formica japonica, Mycetarotes parallelus, Mycocepurus goeldii, Odontomachus brunneus, Sericomyrmex parvulus, Trachymyrmex septentrionalis*

Subset 2: ***Aphaenogaster treatae****, D. quadriceps, F. japonica, M. parallelus, M. goeldii, O. brunneus, S. parvulus, T. septentrionalis*

Subset 3: ***Aphaenogaster ashmeadi****, D. quadriceps, F. japonica, M. parallelus, M. goeldii, O. brunneus, S. parvulus, T. septentrionalis*

Subset 4: *A. floridana,* ***Dinoponera australis****, F. japonica, M. parallelus, M. goeldii, O. brunneus, S. parvulus, T. septentrionalis*

Subset 5: *A. floridana, D. quadriceps,* ***Formica archboldi****, M. parallelus, M. goeldii, O. brunneus, S.* *parvulus, T. septentrionalis*

Subset 6: *A. floridana, D. quadriceps,* ***Formica dolosa****, M. parallelus, M. goeldii, O. brunneus, S.* *parvulus, T. septentrionalis*

Subset 7: *A. floridana, D. quadriceps,* ***Formica pallidefulva****, M. parallelus, M. goeldii, O. brunneus, S.* *parvulus, T. septentrionalis*

Subset 8: *A. floridana, D. quadriceps,* ***Formica subaenescens****, M. parallelus, M. goeldii, O. brunneus, S.* *parvulus, T. septentrionalis*

Subset 9: *A. floridana, D. quadriceps, F. japonica,* ***Mycetarotes acutus****, M. goeldii, O. brunneus, S. parvulus, T. septentrionalis*

Subset 10: *A. floridana, D. quadriceps, F. japonica, M. parallelus,* ***Mycocepurus smithii****, O. brunneus, S. parvulus, T. septentrionalis*

Subset 11: *A. floridana, D. quadriceps, F. japonica, M. parallelus, M. goeldii,* ***Odontomachus chelifer****, S.* *parvulus, T. septentrionalis*

Subset 12: *A. floridana, D. quadriceps, F. japonica, M. parallelus, M. goeldii, O. brunneus,* ***Sericomyrmex amabilis****, T. septentrionalis*

Subset 13: *A. floridana, D. quadriceps, F. japonica, M. parallelus, M. goeldii, O. brunneus,* ***Sericomyrmex mayri****, T. septentrionalis*

Subset 14: *A. floridana, D. quadriceps, F. japonica, M. parallelus, M. goeldii, O. brunneus,* ***Sericomyrmex saramama****, T. septentrionalis*

Subset 15: *A. floridana, D. quadriceps, F. japonica, M. parallelus, M. goeldii, O. brunneus,* ***Sericomyrmex bondari****, T. septentrionalis*

Subset 16: *A. floridana, D. quadriceps, F. japonica, M. parallelus, M. goeldii, O. brunneus, S.* *parvulus,* ***Trachymyrmex holmgreni***

Mean Distance

**Table S5:** Mean Distance ~ Log(Colony Size), PGLS λ= Max. Likelihood (N = 21 genera)

| **Data Subset** | **Coefficient (β)** | **Standard**  **Error** | **AIC** | **LogLik** | **R^2^** | **Lambda (λ)** | **95% CI for λ** | **p-value** |
| --- | --- | --- | --- | --- | --- | --- | --- | --- |
| 1 | 1.364 | 0.248 | 87.215 | -41.607 | 0.615 | 0.586 | NA, NA | **<0.001** |
| 2 | 1.362 | 0.247 | 87.194 | -41.597 | 0.616 | 0.591 | NA, NA | **<0.001** |
| 3 | 1.376 | 0.245 | 87.273 | -41.637 | 0.625 | 0.570 | NA, NA | **<0.001** |
| 4 | 1.075 | 0.299 | 97.793 | -46.897 | 0.404 | 0.545 | NA, NA | **0.002** |
| 5 | 1.347 | 0.245 | 86.589 | -41.294 | 0.615 | 0.603 | NA, NA | **<0.001** |
| 6 | 1.346 | 0.245 | 86.601 | -41.301 | 0.613 | 0.604 | NA, NA | **<0.001** |
| 7 | 1.333 | 0.245 | 86.701 | -41.350 | 0.610 | 0.610 | NA, NA | **<0.001** |
| 8 | 1.349 | 0.246 | 86.722 | -41.361 | 0.613 | 0.598 | NA, NA | **<0.001** |
| 9 | 1.293 | 0.231 | 88.172 | -42.086 | 0.623 | 0.222 | NA, NA | **<0.001** |
| 10 | 1.342 | 0.239 | 87.332 | -41.666 | 0.625 | 0.557 | NA, NA | **<0.001** |
| 11 | 1.421 | 0.249 | 86.947 | -41.474 | 0.631 | 0.525 | NA, NA | **<0.001** |
| 12 | 1.347 | 0.254 | 88.216 | -42.108 | 0.596 | 0.594 | NA, NA | **<0.001** |
| 13 | 1.270 | 0.256 | 89.541 | -42.770 | 0.565 | 0.597 | NA, NA | **<0.001** |
| 14 | 1.345 | 0.231 | 87.252 | -41.626 | 0.640 | 0.283 | NA, NA | **<0.001** |
| 15 | 1.314 | 0.258 | 89.212 | -42.606 | 0.577 | 0.577 | NA, NA | **<0.001** |
| 16 | 1.368 | 0.249 | 87.225 | -41.613 | 0.613 | 0.568 | NA, NA | **<0.001** |

**Table S6:** Mean Distance ~ Log(Colony Size), PGLS λ= 1 (N = 21 genera)

| **Data Subset** | **Coefficient (β)** | **Standard Error** | **AIC** | **LogLik** | **p-value** |
| --- | --- | --- | --- | --- | --- |
| 1 | 1.142 | 0.254 | 89.830 | -41.915 | **<0.001** |
| 2 | 1.142 | 0.252 | 89.807 | -41.903 | **<0.001** |
| 3 | 1.162 | 0.251 | 90.042 | -42.021 | **<0.001** |
| 4 | 0.871 | 0.300 | 98.408 | -46.204 | **<0.001** |
| 5 | 1.129 | 0.250 | 89.236 | -41.618 | **<0.001** |
| 6 | 1.128 | 0.251 | 89.237 | -41.618 | **<0.001** |
| 7 | 1.119 | 0.250 | 89.263 | -41.631 | **<0.001** |
| 8 | 1.129 | 0.251 | 89.350 | -41.675 | **<0.001** |
| 9 | 1.036 | 0.239 | 90.737 | -42.369 | **<0.001** |
| 10 | 1.127 | 0.245 | 89.937 | -41.969 | **<0.001** |
| 11 | 1.173 | 0.256 | 89.886 | -41.943 | **<0.001** |
| 12 | 1.112 | 0.258 | 90.729 | -42.364 | **<0.001** |
| 13 | 1.014 | 0.249 | 91.763 | -42.881 | **<0.001** |
| 14 | 1.065 | 0.242 | 90.616 | -42.308 | **<0.001** |
| 15 | 1.057 | 0.258 | 91.654 | -42.827 | **<0.001** |
| 16 | 1.179 | 0.262 | 90.199 | -42.099 | **<0.001** |

**Table S7:** Mean Distance ~ Log(Colony Size), PGLS λ= 0 (N = 21 genera)

| **Data Subset** | **Coefficient (β)** | **Standard Error** | **AIC** | **LogLik** | **p-value** |
| --- | --- | --- | --- | --- | --- |
| 1 | 1.360 | 0.239 | 91.287 | -42.643 | **<0.001** |
| 2 | 1.360 | 0.239 | 91.279 | -42.640 | **<0.001** |
| 3 | 1.365 | 0.238 | 91.333 | -42.667 | **<0.001** |
| 4 | 1.040 | 0.286 | 100.980 | -47.490 | **0.002** |
| 5 | 1.333 | 0.238 | 90.675 | -42.337 | **<0.001** |
| 6 | 1.333 | 0.238 | 90.683 | -42.342 | **<0.001** |
| 7 | 1.329 | 0.239 | 90.802 | -42.401 | **<0.001** |
| 8 | 1.343 | 0.238 | 90.833 | -42.417 | **<0.001** |
| 9 | 1.290 | 0.227 | 91.414 | -42.707 | **<0.001** |
| 10 | 1.354 | 0.231 | 91.157 | -42.579 | **<0.001** |
| 11 | 1.432 | 0.246 | 90.834 | -42.417 | **<0.001** |
| 12 | 1.349 | 0.248 | 92.345 | -43.173 | **<0.001** |
| 13 | 1.289 | 0.254 | 93.526 | -43.763 | **<0.001** |
| 14 | 1.337 | 0.226 | 90.660 | -42.330 | **<0.001** |
| 15 | 1.323 | 0.253 | 93.224 | -43.612 | **<0.001** |
| 16 | 1.349 | 0.241 | 91.228 | -42.614 | **<0.001** |

Standardized Mean Distance - Chain Networks

**Table S8:** Mean Distance Stdz. to Chains~ Log(Colony Size), PGLS λ= 1 (N = 21 genera)

| **Data Subset** | **Coefficient (β)** | **Standard Error** | **AIC** | **LogLik** | **p-value** |
| --- | --- | --- | --- | --- | --- |
| 1 | -0.055 | 0.024 | -0.514 | 3.257 | **0.031** |
| 2 | -0.056 | 0.024 | -0.337 | 3.168 | **0.028** |
| 3 | -0.055 | 0.023 | -0.513 | 3.257 | **0.028** |
| 4 | -0.059 | 0.019 | -5.645 | 5.823 | **0.007** |
| 5 | -0.051 | 0.022 | -2.465 | 4.233 | **0.034** |
| 6 | -0.050 | 0.024 | 0.406 | 2.797 | 0.054 |
| 7 | -0.051 | 0.022 | -2.380 | 4.190 | **0.035** |
| 8 | -0.053 | 0.023 | -2.234 | 4.117 | **0.031** |
| 9 | -0.049 | 0.022 | -0.093 | 3.047 | **0.036** |
| 10 | -0.052 | 0.021 | -4.107 | 5.053 | **0.021** |
| 11 | -0.054 | 0.024 | -0.521 | 3.261 | **0.035** |
| 12 | -0.053 | 0.024 | -0.311 | 3.156 | **0.035** |
| 13 | -0.052 | 0.022 | -0.332 | 3.166 | **0.028** |
| 14 | -0.050 | 0.022 | -0.027 | 3.014 | **0.039** |
| 15 | -0.050 | 0.023 | 0.066 | 2.967 | **0.042** |
| 16 | -0.057 | 0.024 | -0.688 | 3.344 | **0.029** |

**Table S9:** Mean Distance Stdz. to Chains~ Log(Colony Size), PGLS λ= 0 (N = 21 genera)

| **Data Subset** | **Coefficient (β)** | **Standard Error** | **AIC** | **LogLik** | **p-value** |
| --- | --- | --- | --- | --- | --- |
| 1 | -0.061 | 0.024 | 3.195 | 1.402 | **0.019** |
| 2 | -0.061 | 0.024 | 3.275 | 1.362 | **0.018** |
| 3 | -0.061 | 0.023 | 3.188 | 1.406 | **0.018** |
| 4 | -0.067 | 0.019 | -2.626 | 4.313 | **0.002** |
| 5 | -0.055 | 0.022 | 0.859 | 2.570 | **0.024** |
| 6 | -0.055 | 0.025 | 4.311 | 0.844 | **0.036** |
| 7 | -0.054 | 0.022 | 0.992 | 2.504 | **0.026** |
| 8 | -0.057 | 0.022 | 1.180 | 2.410 | **0.020** |
| 9 | -0.058 | 0.022 | 3.064 | 1.468 | **0.016** |
| 10 | -0.062 | 0.022 | 2.359 | 1.821 | **0.012** |
| 11 | -0.058 | 0.026 | 5.102 | 0.449 | **0.037** |
| 12 | -0.059 | 0.024 | 3.506 | 1.247 | **0.022** |
| 13 | -0.056 | 0.024 | 3.371 | 1.314 | **0.028** |
| 14 | -0.060 | 0.022 | 2.751 | 1.624 | **0.014** |
| 15 | -0.058 | 0.024 | 3.871 | 1.064 | **0.027** |
| 16 | -0.060 | 0.024 | 3.507 | 1.247 | **0.022** |

Standardized Mean Distance – Triangulated Networks

**Table S10:** Mean Distance Stdz. to Triang. ~ Log(Colony Size), OLS of PIC (N = 21 genera)

| **Data Subset** | **Coefficient (β)** | **95% CI for β (0.025,0.975)** | **R^2^** | **p-value** | **AIC** |
| --- | --- | --- | --- | --- | --- |
| 1 | 0.236 | **0.055, 0.466** | 0.348 | 0.005 | -52.202 |
| 2 | 0.237 | **0.070, 0.477** | 0.351 | 0.005 | -52.199 |
| 3 | 0.248 | **0.070, 0.462** | 0.376 | 0.003 | -51.962 |
| 4 | 0.154 | -0.063, 0.396 | 0.138 | 0.097 | -43.149 |
| 5 | 0.239 | **0.076, 0.472** | 0.353 | 0.005 | -52.188 |
| 6 | 0.241 | **0.069, 0.483** | 0.349 | 0.005 | -51.614 |
| 7 | 0.236 | **0.072, 0.477** | 0.349 | 0.005 | -52.213 |
| 8 | 0.238 | **0.061, 0.472** | 0.351 | 0.005 | -52.214 |
| 9 | 0.226 | **0.076, 0.440** | 0.370 | 0.003 | -52.797 |
| 10 | 0.235 | **0.048, 0.452** | 0.363 | 0.004 | -52.246 |
| 11 | 0.248 | **0.081, 0.478** | 0.378 | 0.003 | -53.025 |
| 12 | 0.227 | **0.042, 0.477** | 0.323 | 0.007 | -51.217 |
| 13 | 0.219 | **0.069, 0.452** | 0.340 | 0.006 | -51.843 |
| 14 | 0.237 | **0.085, 0.448** | 0.381 | 0.003 | -52.283 |
| 15 | 0.217 | **0.056, 0.471** | 0.311 | 0.009 | -51.090 |
| 16 | 0.242 | **0.051, 0.492** | 0.319 | 0.008 | -49.470 |

Standardized Mean Distance – MSTs

**Table S11:** Mean Distance Stdz. to MSTs ~ Log(Colony Size), OLS of PIC (N = 21 genera)

| **Data Subset** | **Coefficient (β)** | **95% CI for β (0.025,0.975)** | **R^2^** | **p-value** | **AIC** |
| --- | --- | --- | --- | --- | --- |
| 1 | 0.110 | **0.022, 0.240** | 0.281 | 0.013 | -76.398 |
| 2 | 0.110 | **0.014, 0.244** | 0.280 | 0.014 | -76.422 |
| 3 | 0.115 | **0.023, 0.244** | 0.303 | 0.010 | -76.241 |
| 4 | 0.065 | -0.055, 0.201 | 0.089 | 0.189 | -67.686 |
| 5 | 0.113 | **0.020, 0.239** | 0.290 | 0.012 | -76.417 |
| 6 | 0.114 | **0.022, 0.251** | 0.283 | 0.013 | -75.448 |
| 7 | 0.111 | **0.018, 0.238** | 0.287 | 0.012 | -76.545 |
| 8 | 0.112 | **0.016, 0.237** | 0.286 | 0.012 | -76.533 |
| 9 | 0.100 | **0.018, 0.212** | 0.271 | 0.016 | -76.112 |
| 10 | 0.112 | **0.009, 0.224** | 0.307 | 0.009 | -76.947 |
| 11 | 0.116 | **0.026, 0.241** | 0.310 | 0.009 | -77.286 |
| 12 | 0.107 | **0.017, 0.234** | 0.265 | 0.017 | -75.884 |
| 13 | 0.097 | **0.023, 0.207** | 0.251 | 0.021 | -75.640 |
| 14 | 0.103 | **0.018, 0.214** | 0.275 | 0.015 | -76.147 |
| 15 | 0.101 | **0.017, 0.242** | 0.251 | 0.021 | -75.582 |
| 16 | 0.113 | **0.012, 0.238** | 0.268 | 0.016 | -74.926 |

Number of Cycles

**Table S12:** Number of Cycles ~ Log(Colony Size), OLS of PIC (N = 21 genera)

| **Data Subset** | **Coefficient (β)** | **95% CI for β (0.025,0.975)** | **R^2^** | **p-value** | **AIC** |
| --- | --- | --- | --- | --- | --- |
| 1 | 0.293 | -0.089, 0.528 | 0.160 | 0.072 | -23.053 |
| 2 | 0.363 | -0.114, 0.626 | 0.222 | 0.031 | -22.278 |
| 3 | 0.357 | -0.057, 0.621 | 0.222 | 0.031 | -22.495 |
| 4 | 0.312 | **0.047, 0.564** | 0.289 | 0.012 | -33.332 |
| 5 | 0.237 | -0.117, 0.478 | 0.105 | 0.152 | -21.775 |
| 6 | 0.234 | -0.162, 0.495 | 0.076 | 0.228 | -15.092 |
| 7 | 0.257 | -0.083, 0.486 | 0.159 | 0.073 | -28.014 |
| 8 | 0.263 | -0.074, 0.489 | 0.164 | 0.068 | -28.076 |
| 9 | 0.262 | -0.008, 0.482 | 0.150 | 0.083 | -22.804 |
| 10 | 0.302 | -0.043, 0.533 | 0.183 | 0.053 | -23.611 |
| 11 | 0.315 | -0.044, 0.597 | 0.152 | 0.081 | -19.023 |
| 12 | 0.286 | -0.056, 0.562 | 0.155 | 0.077 | -22.932 |
| 13 | 0.248 | -0.066, 0.504 | 0.132 | 0.105 | -22.391 |
| 14 | 0.264 | -0.021, 0.533 | 0.148 | 0.085 | -22.764 |
| 15 | 0.271 | -0.055, 0.523 | 0.146 | 0.088 | -22.708 |
| 16 | 0.303 | -0.058, 0.546 | 0.165 | 0.068 | -23.158 |

Number of Cycles Standardized to Triangulated Networks

**Table S13:** Standardized Number of Cycles ~ Log(Colony Size), OLS of PIC (N = 21 genera)

| **Data Subset** | **Coefficient (β)** | **95% CI for β (0.025,0.975)** | **R^2^** | **p-value** | **AIC** |
| --- | --- | --- | --- | --- | --- |
| 1 | -0.048 | -0.102, -0.015 | 0.090 | 0.186 | -82.571 |
| 2 | -0.030 | -0.093, -0.006 | 0.038 | 0.396 | -82.394 |
| 3 | -0.031 | -0.098, -0.009 | 0.042 | 0.373 | -82.506 |
| 4 | -0.040 | -0.089, -0.009 | 0.072 | 0.241 | -82.172 |
| 5 | -0.049 | -0.108, -0.015 | 0.094 | 0.176 | -82.585 |
| 6 | -0.049 | -0.106, -0.017 | 0.094 | 0.177 | -82.525 |
| 7 | -0.048 | -0.101, -0.016 | 0.092 | 0.182 | -82.613 |
| 8 | -0.048 | -0.105, -0.016 | 0.092 | 0.181 | -82.621 |
| 9 | -0.081 | -0.184, -0.021 | 0.249 | 0.021 | -82.229 |
| 10 | -0.037 | -0.086, -0.004 | 0.060 | 0.285 | -81.918 |
| 11 | -0.049 | -0.105, -0.016 | 0.095 | 0.174 | -82.723 |
| 12 | -0.049 | -0.117, -0.014 | 0.098 | 0.166 | -82.755 |
| 13 | -0.048 | -0.121, -0.009 | 0.105 | 0.152 | -82.906 |
| 14 | -0.108 | -0.246, 0.009 | 0.202 | 0.041 | -66.168 |
| 15 | -0.050 | -0.116, -0.013 | 0.103 | 0.155 | -82.865 |
| 16 | -0.052 | -0.114, -0.015 | 0.103 | 0.156 | -82.855 |

**PHYLOGENETIC SIGNAL**

Species subsets refer to the sets listed for each trait above for Tables S3-13. Colony size contains the same species subsets as Chamber Number.

**Table S14:** Colony Size - Phylogenetic Signal (N = 24 genera)

| **Data Subset** | **Est. Pagel’s λ** | **p-value** | **AIC Est. λ Model** | **AIC BM Model** | **AIC No Sig Model** |
| --- | --- | --- | --- | --- | --- |
| 1 | 0.440 | 0.348 | 99.034 | 99.353 | 97.286 |
| 2 | 0.450 | 0.340 | 98.965 | 99.146 | 97.248 |
| 3 | 0.427 | 0.364 | 99.164 | 99.537 | 97.359 |
| 4 | 0.396 | 0.405 | 99.487 | 99.990 | 97.554 |
| 5 | 0.519 | 0.186 | 101.455 | 101.440 | 100.575 |
| 6 | 0.391 | 0.471 | 100.757 | 100.909 | 98.648 |
| 7 | 0.147 | 1.000 | 98.244 | 99.358 | 95.616 |
| 8 | 0.305 | 0.576 | 102.011 | 102.916 | 99.696 |
| 9 | 0.593 | 0.241 | 98.176 | 97.067 | 96.919 |
| 10 | 0.528 | 0.282 | 98.439 | 97.837 | 96.970 |
| 11 | 0.624 | 0.225 | 98.564 | 97.393 | 97.406 |
| 12 | 0.625 | 0.223 | 98.262 | 96.993 | 97.115 |
| 13 | 0.541 | 0.292 | 100.347 | 100.271 | 98.827 |
| 14 | 0.474 | 0.320 | 98.777 | 98.707 | 97.136 |
| 15 | 0.421 | 0.374 | 99.486 | 100.749 | 97.646 |
| 16 | 0.407 | 0.406 | 98.740 | 99.173 | 96.802 |
| 17 | 0.412 | 0.396 | 98.708 | 99.137 | 96.801 |
| 18 | 0.400 | 0.419 | 98.812 | 99.244 | 96.835 |
| 19 | 0.431 | 0.361 | 98.831 | 99.201 | 97.036 |

**Table S15:** Chamber Number - Phylogenetic Signal (N = 24 genera)

| **Data Subset** | **Est. Pagel’s λ** | **p-value** | **AIC Est. λ Model** | **AIC BM Model** | **AIC No Sig Model** |
| --- | --- | --- | --- | --- | --- |
| 1 | 1.066 | 0.091 | 182.720 | 180.092 | 182.598 |
| 2 | 1.091 | 0.049 | 182.348 | 179.719 | 182.702 |
| 3 | 1.066 | 0.116 | 183.277 | 180.649 | 182.791 |
| 4 | 1.066 | 0.162 | 184.090 | 181.461 | 183.108 |
| 5 | 1.066 | 0.156 | 183.540 | 180.911 | 182.603 |
| 6 | 1.066 | 0.088 | 182.776 | 180.148 | 182.691 |
| 7 | 1.085 | 0.119 | 191.689 | 189.060 | 190.660 |
| 8 | 1.066 | 0.091 | 182.737 | 180.108 | 182.612 |
| 9 | 1.063 | 0.066 | 183.119 | 180.491 | 183.549 |
| 10 | 1.066 | 0.062 | 182.878 | 180.249 | 183.355 |
| 11 | 1.060 | 0.070 | 183.263 | 180.634 | 183.652 |
| 12 | 1.068 | 0.060 | 182.725 | 180.096 | 183.207 |
| 13 | 1.060 | 0.070 | 183.263 | 180.634 | 183.652 |
| 14 | 1.070 | 0.062 | 182.550 | 179.922 | 182.949 |
| 15 | 1.027 | 0.133 | 183.311 | 180.682 | 182.898 |
| 16 | 1.063 | 0.097 | 181.168 | 178.540 | 180.972 |
| 17 | 1.063 | 0.094 | 181.318 | 178.690 | 181.159 |
| 18 | 1.063 | 0.094 | 181.318 | 178.690 | 181.159 |
| 19 | 1.064 | 0.092 | 182.003 | 179.374 | 181.878 |

**Table S16:** Standardized Chamber Width – Phylogenetic Signal (N = 21 genera)

| **Data Subset** | **Est. Pagel’s λ** | **p-value** | **AIC Est. λ Model** | **AIC BM Model** | **AIC No Sig Model** |
| --- | --- | --- | --- | --- | --- |
| 1 | 0.712 | 0.084 | 88.785 | 86.834 | 89.023 |
| 2 | 0.804 | **0.040** | 88.426 | 86.508 | 89.885 |
| 3 | 0.718 | 0.076 | 88.321 | 86.382 | 88.716 |
| 4 | 0.725 | 0.068 | 87.787 | 85.865 | 88.376 |
| 5 | 0.684 | 0.094 | 88.953 | 87.097 | 89.005 |
| 6 | 1.123 | **0.019** | 86.619 | 83.874 | 87.562 |
| 7 | 0.681 | 0.088 | 89.090 | 87.348 | 89.256 |
| 8 | 1.123 | **0.011** | 85.793 | 83.048 | 86.919 |
| 9 | 0.746 | 0.080 | 88.484 | 86.340 | 88.796 |
| 10 | 0.425 | 0.158 | 93.925 | 95.910 | 93.171 |
| 11 | 0.314 | 0.245 | 98.371 | 103.399 | 96.979 |
| 12 | 0.772 | 0.068 | 88.479 | 86.273 | 89.071 |
| 13 | 0.708 | 0.082 | 88.380 | 86.473 | 88.653 |
| 14 | 0.715 | 0.085 | 89.066 | 87.084 | 89.279 |
| 15 | 0.711 | 0.084 | 88.729 | 86.784 | 88.972 |

**Table S17:** Mean Distance Stdz. to Chain - Phylogenetic Signal (N = 21 genera)

| **Data Subset** | **Est. Pagel’s λ** | **p-value** | **AIC Est. λ Model** | **AIC BM Model** | **AIC No Sig Model** |
| --- | --- | --- | --- | --- | --- |
| 1 | 1.124 | **0.001** | -2.222 | -4.967 | -2.194 |
| 2 | 1.124 | **0.002** | -1.805 | -4.551 | -2.031 |
| 3 | 1.124 | **0.001** | -2.021 | -4.766 | -2.117 |
| 4 | 1.124 | **0.000** | -5.465 | -8.210 | -2.287 |
| 5 | 1.124 | **0.001** | -4.963 | -7.708 | -4.345 |
| 6 | 1.124 | **0.001** | -2.222 | -4.967 | -2.194 |
| 7 | 1.124 | **0.001** | -2.222 | -4.967 | -2.194 |
| 8 | 1.124 | **0.003** | -1.943 | -4.688 | -2.798 |
| 9 | 1.124 | **0.001** | -2.222 | -4.967 | -2.194 |
| 10 | 1.124 | **0.001** | -2.222 | -4.967 | -2.194 |
| 11 | 1.124 | **0.001** | -2.222 | -4.967 | -2.194 |
| 12 | 1.124 | **0.002** | -4.515 | -7.261 | -5.276 |
| 13 | 1.124 | **0.001** | -2.234 | -4.979 | -2.209 |
| 14 | 1.124 | **0.002** | -4.505 | -7.250 | -5.249 |
| 15 | 1.124 | **0.001** | -2.417 | -5.162 | -1.327 |
| 16 | 1.124 | **0.002** | -4.109 | -6.854 | -4.561 |

**Table S18:** Number of Cycles – Phylogenetic Signal (N = 21 genera)

| **Data Subset** | **Est. Pagel’s λ** | **p-value** | **AIC Est. λ Model** | **AIC BM Model** | **AIC No Sig Model** |
| --- | --- | --- | --- | --- | --- |
| 1 | 1.124 | **0.005** | 74.979 | 72.234 | 72.387 |
| 2 | 0.000 | 1.000 | 75.366 | 74.654 | 72.620 |
| 3 | 0.000 | 1.000 | 75.281 | 74.416 | 72.536 |
| 4 | 1.124 | **0.005** | 74.979 | 72.234 | 72.387 |
| 5 | 1.124 | **0.000** | 67.679 | 64.934 | 69.365 |
| 6 | 1.124 | **0.005** | 74.979 | 72.234 | 72.387 |
| 7 | 1.124 | **0.005** | 74.979 | 72.234 | 72.387 |
| 8 | 1.124 | **0.005** | 74.979 | 72.234 | 72.387 |
| 9 | 1.124 | **0.005** | 74.979 | 72.234 | 72.387 |
| 10 | 1.124 | **0.005** | 74.979 | 72.234 | 72.387 |
| 11 | 1.124 | **0.005** | 74.979 | 72.234 | 72.387 |
| 12 | 1.124 | **0.005** | 74.979 | 72.234 | 72.387 |
| 13 | 0.000 | 1.000 | 81.318 | 78.573 | 78.689 |
| 14 | 1.124 | **0.004** | 69.736 | 66.991 | 67.636 |
| 15 | 1.124 | **0.002** | 78.996 | 76.251 | 77.574 |
| 16 | 1.124 | **0.004** | 69.804 | 67.059 | 68.051 |

**R CODE - GENERATING REFERENCE NETWORKS**

```
library(igraph)
library(deldir)
library(mcMST)
library(ggplot2)

#Generating Reference Models
#PLANAR & MST ----
#Use to generate random x-y coords for planar networks
# n=# nodes#

# initiate variable to fill in loops
error.list<-c()

mean.cycles.triang<-c()
se.cycles.triang<-c()

mean.md.planar<-c()
se.md.planar<-c()

mean.md.mst<-c()
se.md.mst<-c()

for (i in (3:100)) { # loop on nest sizes from 3 to 100 chambers

  cycles.p<-c()
  md.p<-c() # initiate variable to collect mean distances for planar
  error.count=0

  md.m<-c() # initiate variable to collect mean distances for MSTs

  for (j in (1:1000)) { # generate 1000 random samples for each nest size

    #build random, spatial set of nodes
    yoda=0
    while (yoda==0){
      x<-sample(i) # generate random x coordinates for i number of nodes
      y<-sample(i) # generate random y coordinates for i number of nodes

      dxy<-deldir(x,y)

      if (!is.null(dxy)) {
        yoda = 1
      } else {
        message('Retried deldir for i = ', i)
        error.count=error.count+1
      }
    }

    vert <- data.frame(
      id1 = dxy$delsgs$ind1,
      id2 = dxy$delsgs$ind2)
    edgelist.planar<-as.matrix(vert) #convert data to a matrix - 2 columns

    #Convert nodes into a triangulated graph
    planar_graph<-graph_from_edgelist(edgelist.planar,directed=FALSE)

    #Convert triangulated graph into a MST
    mintree<-mst(planar_graph)
    #plot(mintree)
    #plot(planar_graph)

    #PLANAR - Calculate Mean Distance &  Num Cycles
    #Mesh
    n.top<-vcount(planar_graph) #number of nodes or vertices
    m.top<-ecount(planar_graph) #number of edges
    cycles.p[j]<-m.top-n.top+1

    #Mean Distance
    #Mean distance = 1/(nodes*(nodes+1))*sum(all pairwise distances)
    md.p[j]<-mean_distance(planar_graph)
    #generates lists of 1000 long for one i value

    #MST - Calculate Mean Distance
    #Mean distance = 1/(nodes*(nodes+1))*sum(all pairwise distances)
    md.m[j]<-mean_distance(mintree)

    #generates lists of 1000 long for one i value

  }

  error.list[i]<-error.count

  #Cycles - TRIANGULATED
  mean.cycles.triang[i]<-mean(cycles.p)
  se.cycles.triang[i]<-sd(cycles.p)/sqrt(length(cycles.p))

  #mean distance - TRIANGULATED
  mean.md.planar[i]<-mean(md.p)
  se.md.planar[i]<-sd(md.p)

  #mean distance - MST
  mean.md.mst[i]<-mean(md.m)
  se.md.mst[i]<-sd(md.m)
}

#END OF GENERATING NULL NETWORKS - TRIANGULATION & MINIMUM SPANNING TREES

#CHAIN----
#GENERATE A CHAIN GRAPH

md.c<-c() # initiate variable to collect mean distances for chain

for (i in 3:100) { # loop on nest sizes from 3 to 100 chambers

  col1<-seq(1:(i-1))
  col2<-seq(from=2, to =i)

  edge.list.chain<-cbind(col1,col2)
  edgelist<-as.matrix(edge.list.chain)
  chain<-graph_from_edgelist(edgelist,directed=FALSE)

  #Mean Distance
  md.c[i]<-mean_distance(chain)

}

#PLOTTING Mean Distance -----

#MEAN DISTANCE
#**PLANAR----
mean.md.planar.data<-as.data.frame(mean.md.planar)
#Add index
#NumNodes <- as.data.frame(seq.int(length(mean.rmd.planar))) # Add index column
#Add SE
SE<-as.data.frame(se.md.planar)
#CI
LoCI<- mean.md.planar.data[,2]-1.96*SE[,2]
UpCI <- mean.md.planar.data[,2]+1.96*SE[,2]
#Combine
mdplan<-cbind(mean.md.planar.data,LoCI,UpCI)
colnames(mdplan)<-c("NumNodes","MD_Planar", "pLo","pUp")

#**MST----
mean.md.mst.data<-as.data.frame(mean.md.mst)
#Add SE
SE<-as.data.frame(se.md.mst)
##CI
LoCI<- mean.md.mst.data[,2]-1.96*SE[,2]
UpCI <- mean.md.mst.data[,2]+1.96*SE[,2]
#Combine
mdmst<-cbind(mean.md.mst.data[,2],LoCI,UpCI)
colnames(mdmst)<-c("MD_MST","mLo","mUp")

#**CHAIN----
newformat1<-cbind(mdplan, mdmst)
colnames(md.c)<-c("NumNode","MD_Chain")
newformat<-cbind(newformat1,md.c[2]) #Added chain network mean distance measures

#Need to add observed data using merge and then add to ggplot at geom_point

#**Observed----
#Observed Nest Data
path = SET TO PATH WHERE FILE SITS
dall<-read.csv(paste (‘path’, ‘\\Trait_Data_WholeNest_2021.csv’))
obs<-dall[,c("Name","mean.dist.all","num.node.pruned")]

#Remove polydomous nests:
#POLYDOMOUS REMOVED----
dall.s<-subset(dall,filename.list!="D_indicum12.csv" & filename.list!="A_balzani_T1.csv" & filename.list!="A_balzani_T2.csv" & filename.list!="A_balzani_T3.csv" & filename.list!="A_balzani_T5.csv" & filename.list!="A_balzani_T6.csv" & filename.list!="A_balzani_T7.csv" & filename.list!="A_balzani_T8.csv" & filename.list!="A_balzani2.csv" & filename.list!="A_balzani6.csv"  & filename.list!="F_japonica_T4.csv")
obs<-dall.s[,c("Name","mean.dist.all","num.node.pruned")]

colnames(obs)<-c("Name","observed", "NumNodes")

newall<-merge(newformat, obs, by="NumNodes" ,all=TRUE)

max(newall$observed,na.rm=TRUE) #Maximum Y-value = 11.99

#*Plot Mean Distance Ref Models and Observed----
ggplot(data = newall) +
  geom_count(aes(x=NumNodes,y=observed),color="grey35")+ #observed
  geom_ribbon(aes(x = NumNodes, ymin = pLo, ymax = pUp),
              fill = "blue3", alpha = 0.3) +
  geom_line(aes(x = NumNodes, y = MD_Planar), linetype="solid",color = "blue3") + #triangulated
  geom_ribbon(aes(x = NumNodes, ymin = mLo, ymax = mUp),
              fill = "goldenrod3", alpha = 0.3) +
  geom_line(aes(x = NumNodes, y = MD_MST), linetype="solid",color = "goldenrod3")+
  geom_line(aes(x = NumNodes, y = MD_Chain), linetype="solid",color = "firebrick4") +
  ggtitle("Mean Distance")+
  xlim(0,101)+
  xlab("Number of Nodes")+
  ylab("Mean Distance")+
  theme_bw() +
  theme(panel.border = element_blank(), panel.grid.major = element_blank(),
        panel.grid.minor = element_blank(),
        axis.line = element_line(colour = "black"),plot.title=element_text(hjust=0.5))+
  scale_y_continuous(limits=c(0,12.5))+
  theme(text = element_text(size=23))

mean.cycles.triang<-read.csv("D:\\Documents\\Research Fun\\Nest Architecture\\DATA & ANALYSES\\Nulls\\Reference Models\\Triangulated CycleNum Meancsv")
se.cycles.triang<-read.csv("D:\\Documents\\Research Fun\\Nest Architecture\\DATA & ANALYSES\\Nulls\\Reference Models\\Triangulated CycleNum SE.csv")

#*Plot Number of cycles in Triangulated Model and Observed----
#*
triang.cycles.data<-as.data.frame(mean.cycles.triang)
SE<-as.data.frame(se.cycles.triang)
##CI
LoCI<- mean.cycles.triang[,2]-1.96*SE[,2]
UpCI <- mean.cycles.triang[,2]+1.96*SE[,2]
#Combine
triang.cycles.data<-cbind(mean.cycles.triang,LoCI,UpCI)
colnames(triang.cycles.data)<-c("NumNodes","Cyc_Triang","Lo","Up")

#Preparing Observed Data----
dall<-
paste (‘path’, ‘\\Trait_Data_WholeNest_2021.csv’)

#Remove polydomous nests:
#POLYDOMOUS REMOVED----
dall.s<-subset(dall,filename.list!="D_indicum12.csv" & filename.list!="A_balzani_T1.csv" & filename.list!="A_balzani_T2.csv" & filename.list!="A_balzani_T3.csv" & filename.list!="A_balzani_T5.csv" & filename.list!="A_balzani_T6.csv" & filename.list!="A_balzani_T7.csv" & filename.list!="A_balzani_T8.csv" & filename.list!="A_balzani2.csv" & filename.list!="A_balzani6.csv" & filename.list!="F_japonica_T4.csv")

obs<-dall.s[,c("Name","cycles.list","num.node.pruned")]
colnames(obs)<-c("Name","observed", "NumNodes")
cycles.all<-merge(triang.cycles.data, obs, by="NumNodes" ,all=TRUE)
max(cycles.all$observed,na.rm=TRUE) #Maximum Y-value = 8

ggplot(data = cycles.all) +
  geom_count(aes(x=NumNodes,y=observed),color="grey35")+ #observed
  geom_line(aes(x = NumNodes, y = Cyc_Triang), linetype="solid",color = "blue3") +
  geom_hline(yintercept = 0,linetype="solid",color="darkorange2")+
  ggtitle("Number of Cycles")+
  xlim(0,101)+
  xlab("Number of Nodes")+
  ylab("Nuber of Cycles in Network")+
  theme_bw() +
  theme(panel.border = element_blank(), panel.grid.major = element_blank(),  text = element_text(size=23),
        panel.grid.minor = element_blank(),
        axis.line = element_line(colour = "black"),plot.title=element_text(hjust=0.5))+
  scale_y_continuous(limits=c(0,150))
```
